# Selective antibiofilm properties and biocompatibility of nano-ZnO and nano-ZnO/Ag coated surfaces

**DOI:** 10.1101/2020.03.18.996967

**Authors:** M. Rosenberg, M. Visnapuu, H. Vija, V. Kisand, K. Kasemets, A. Kahru, A. Ivask

## Abstract

Spread of pathogenic microbes and antibiotic-resistant bacteria in health-care settings and public spaces is a serious public health challenge. Materials that prevent solid surface colonization or impede touch-transfer of viable microbes could provide means to decrease pathogen transfer from high-touch surfaces in critical applications. ZnO and Ag nanoparticles have shown great potential in antimicrobial applications. Less is known about nano-enabled surfaces. Here we demonstrate that surfaces coated with nano-ZnO or nano-ZnO/Ag composites are not cytotoxic to human keratinocytes and possess species-selective medium-dependent antibiofilm activity against *Escherichia coli, Staphylococcus aureus* and *Candida albicans*. Colonization of nano-ZnO and nano-ZnO/Ag surfaces by *E. coli* and *S. aureus* was decreased in static oligotrophic conditions (no planktonic growth). Moderate to no effect was observed for bacterial biofilms in growth medium (supporting exponential growth). Inversely, nano-ZnO surfaces enhanced biofilm formation by *C. albicans* in oligotrophic conditions. However, enhanced *C. albicans* biofilm formation on nano-ZnO surfaces was effectively counteracted by the addition of Ag. Possible selective enhancement of biofilm formation by the yeast *C. albicans* on Zn-enabled surfaces should be taken into account in antimicrobial surface development. Our results also indicated the importance of the use of application-appropriate test conditions and exposure medium in antimicrobial surface testing.

## INTRODUCTION

Biofilms are by far the preferred lifestyle of bacteria [1], mostly in diverse nutrient-limited environmental niches. Biofilm communities cause biomass buildup on solid surfaces that results in major expenses in marine traffic, water systems maintenance and in the industrial sector. Biofilms can also harbor potential human pathogens in food industry, health-care facilities, drinking water systems and on high-touch surfaces in public spaces. It is estimated that hard to treat pathogenic biofilms account for over 80% of all human microbial infections [2] with antibiotic-resistant ESKAPE pathogens [3] including six nosocomial pathogens commonly associated with multidrug resistance and virulence (*Enterococcus faecium, Staphylococcus aureus, Klebsiella pneumoniae, Acinetobacter baumannii, Pseudomonas aeruginosa, Enterobacter spp*.) being the most problematic in the field. Biofilms pose a major medical challenge as they can be over 1000-fold more tolerant to antibiotics than their planktonic counterparts [4]. Spatially heterogeneous natural selection in biofilm milieu also contributes to antibiotic resistance development and transfer [5], [6] as well as producing antibiotic-resistant bacteria that are more fit and not easily outcompeted in the absence of the drug [7].

Controversially, most of the methods used to assess antimicrobial properties of surface materials, irrespective of their proposed application, either use planktonic cultures or study indirect effects such as release of antimicrobial compounds from the surfaces [8]. Such methods might not correctly report surface efficacy in proposed applications. Even if biofilm formation on material of interest is studied, often-used methods of biofilm viability assessment are prone to critical failures. For example, viability staining with propidium iodide and suitable counterstains can dramatically underestimate viability in both oligotrophic [9] and growth medium biofilms [10] while colony counts, the gold standard of microbiology, depend on viable cell harvesting efficiency and dispersion of harvested aggregates prior to cultivation [11], [12].

Susceptibility to metal ions has been demonstrated to be similar for both biofilms and planktonic cultures [13] which makes antimicrobial metal-based approaches good candidates to prevent pathogen transfer and biofilm formation on high-touch surfaces. Silver and copper are good examples of infection prevention and marine antifouling agents historically used even before deeper knowledge of microbes or biofilms was established. Historical use of the former is well reviewed by Lemire *et al*. [14]. Zinc is a later addition to the list after wider use of ZnO nanoparticles. Although zinc is an essential micronutrient being incorporated into 4-10% of proteins across the domains of life [15], it possesses a dose-dependent antibacterial activity at higher concentrations. Zinc toxicity towards microbes is mainly attributable to deactivation of proteins via thiol-disulfide chemistry [14], [16] and protein binding or metal replacement (e.g. manganese starvation[17], [18]) resulting in impaired energy metabolism [19]–[21], higher susceptibility to reactive oxygen species (ROS) [22] and eventually loss of membrane potential and membrane permeabilization. Zn tolerance in non-physiological concentrations has been shown to depend on microbial species with *C. albicans* and *P. aeruginosa* being less sensitive to Zn toxicity than *E. coli* or *S. aureus* [23]. ZnO nanoparticles have been demonstrated to possess additional antibacterial properties due to direct contact and ROS production [24]. Nano-specific effects damage bacterial cell membranes and downregulate genes associated with managing oxidative stress in *S. aureus* as well as upregulate genes associated with cation efflux in *E. coli* and *P. aeruginosa* and inhibit biofilm formation by *E. coli, S. aureus* and *P. aeruginosa* [25]–[30] among other microbes. Addition of silver adds to antimicrobial effects in a variety of mechanisms including membrane and cell wall damage, protein inactivation, impaired energy metabolism, ROS production and DNA damage [31]–[39].

Inhibition of biofilm formation on Zn-based applications can partially be attributed to general Zn toxicity to bacteria above physiological concentrations but also other biofilm-specific mechanisms of action could be involved. For example, it has been proposed that sublethal Zn or Ag concentrations could affect biofilm formation by interfering with quorum sensing [40]–[43] or modulate amyloid fibril formation [44], [45].

Another promising way is the development of antimicrobial surfaces containing photocatalysts that not only induce microbial killing but also degradation of organic matter under specific illumination conditions that could hopefully decrease dry touch-transfer of pathogens or delay moist surface colonization between light exposures. The most popular photocatalyst is TiO_2_ but also ZnO is widely used [24]. In one of our previous studies [46] we demonstrated that in addition to killing bacterial cells during photocatalysis, TiO_2_ surfaces also caused photooxidation of bacterial debris thus, referring to the possibility of extended efficacy of those surfaces.

The idea behind photodegradation is that light with high enough energy to exceed band gap energy of the photocatalytic metal oxide (e.g. 3.37 eV in the case of ZnO) will excite electron-hole pairs. The photogenerated electrons (e^-^) and holes (h^+^) can reduce or oxidize compounds like surface adsorbed O_2_ and H_2_O to produce ROS. The produced ROS (e.g. superoxide anion radical ^•^O_2_^-^ and hydroxyl radical ^•^OH) are able to partially or completely degrade organic contaminants including microbes. It has been claimed that ZnO can produce ROS even without light activation [47] - ROS production via releasing of trapped electrons, so called “photocatalysis in the dark” [48] with respective antimicrobial activity [49], [50]. Therefore, not only metal ions but assumingly also ROS can contribute to the antimicrobial and self-cleaning nature of photocatalytic metal oxide based surfaces in dark conditions. Increased antimicrobial and antibiofilm behavior of nano-ZnO covered dental implant [51] and ZnO thin film covered food-packaging polymer [52] compared to uncoated controls indicate a promising potential of ZnO-based coatings. The use of nanoparticle-based surfaces increases the potential efficiency and reactivity of such surfaces due to increased specific surface area of nanosized matter.

We have previously reported developing novel multi-effective antimicrobial coatings based on nano-ZnO/Ag composite particles [53]. The novelty of these coatings rises from a combined and complex effect of different antimicrobial mechanisms: (i) antimicrobial activity of Zn^2+^ ions, (ii) antimicrobial activity of Ag^+^ ions, and (iii) antimicrobial activity of ROS, generated at the surface of nano-ZnO and nano-ZnO/Ag via photocatalytic processes under UV-A illumination. Our coatings also have two additional advantages (i) degradation of organic debris (incl. dead bacteria) by ROS takes place on the surface and (ii) photocatalytic activity of ZnO is enhanced by the formation of nano-ZnO/Ag composite particles (via charge separation process in ZnO/Ag system).

Antimicrobial properties of our previously developed UV-A-induced nano-ZnO and nano-ZnO/Ag composite coated surfaces were evaluated using an in-house protocol based on ISO standards for measurement of antibacterial activity of non-porous surfaces (ISO 22196:2011 [54]) and photocatalytic surfaces (ISO 27447:2009 [55]), protocols designed to measure antimicrobial action in a thin layer of microbial suspension uniformly spread between test surface and cover film. The nano-ZnO-coated solid surfaces were found to be highly effective under UV-A illumination with over 3 log decrease in planktonic *E. coli* and *S. aureus* viability during 1 h exposure. Photodepositing Ag onto the nano-ZnO increased its photocatalytic activity and acted as an additional antimicrobial agent in the absence of UV-A exposure. [53]

With the intent to further develop these materials for use on high-touch surfaces in the public spaces, in this study we additionally assessed the efficacy of the surface materials against biofilm formation in application-appropriate oligotrophic environment and in the absence of UV-A exposure. For that we opted for a static model to grow biofilms directly on the studied surfaces and to be able to simultaneously monitor dissolution-driven toxicity to the planktonic cells in a small closed system. A mixed approach combining physical methods (vortexing and ultrasonication) with chemical ones (addition of surfactant, using high salt concentration) was used for biofilm harvesting. Biocompatibility of the surfaces was studied using *in vitro* skin-relevant human cell growth directly on the nano-enabled surfaces.

## RESULTS

To assess both anti-adhesion effect of the coating materials and metal release-associated antimicrobial activity towards planktonic microbes, biofilm formation on the sparsely and densely coated nano-ZnO and sparsely coated nano-ZnO/Ag surfaces as well as viability of planktonic microbes in the test system was studied. Biofilms were either grown in static oligotrophic environment (1:500 diluted nutrient broth in synthetic tap water, not supporting planktonic growth) at room temperature to mimic real-life-like use in moist environments (similar to previously used standard conditions for planktonic testing [53]) or in nutrient-rich growth mediums (LB, YPD; supporting exponential growth) resembling classical laboratory approach for studying biofilms [56]. Well described human-relevant biofilm-forming Gram-positive (*S. aureus* ATCC25923) and Gram-negative (*E. coli* MG1655) model bacteria were selected for the experiments. *C. albicans* CAI4 was included as a fungal model organism to represent different types of microbes potentially transferred by fomites. The effect of the surfaces on biofilm formation was quantitively evaluated by harvesting viable adherent cells followed by colony counting or by qualitative epifluorescence microscopy. In parallel, antimicrobial activity towards planktonic cells above the surfaces was analyzed using colony counting.

### Biofilm formation on surfaces coated with nano-ZnO or nano-ZnO/Ag

Nano-ZnO inhibited bacterial biofilm formation in a dose-dependent manner in oligotrophic conditions (Fig. 1a, 1b) reaching maximum of 2.12 and 3.49 log reduction on dense nano-ZnO surface compared to uncoated surface after 72 h for *E. coli* and *S. aureus*, respectively. In growth medium, however, only the dense nano-ZnO surface significantly inhibited biofilm formation of *S. aureus* but not *E. coli* (Fig. 1d, 1e). It was also evident that while there was rapid surface colonization taking place during the first 3 h of incubation in oligotrophic conditions, regardless of surface type, the surfaces in growth medium were either colonized more slowly or the initial adherence was weaker and adherent cells were more easily washed off during sample preparation.

**Figure 1.**
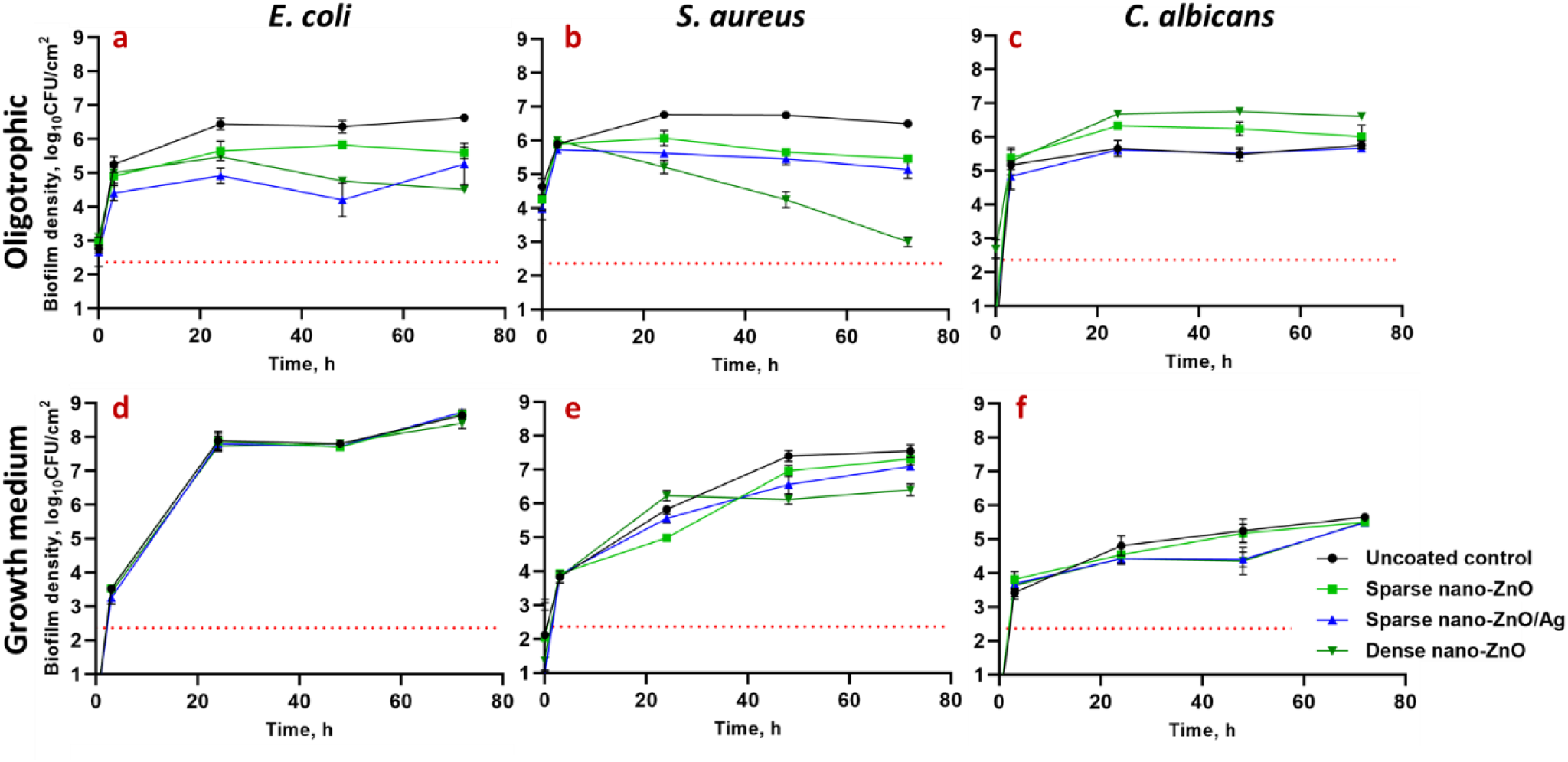
*E. coli* MG1655 (a, d), *S. aureus* ATCC25923 (b, e) or *C. albicans* CAI4 (c, f) monospecies biofilms harvested at different time points from nano-ZnO or nano-ZnO/Ag or uncoated surfaces in static oligotrophic environment (1:500 diluted NB in synthetic tap water; a, b, c) or growth medium (LB: d, e; YPD: f). Biofilm formation was more affected by the nano-enabled coatings in oligotrophic environment than in growth medium with significantly decreased colonization by *E. coli* and *S. aureus* from 24 h onwards while nano-ZnO surfaces selectively enhanced surface colonization by *C. albicans*. Only dense nano-ZnO surfaces demonstrated statistically significant moderate colonization inhibiting effect on *S. aureus* in growth medium 48-72 h post-inoculation. Red dotted line represents the limit of quantification (231 CFU/cm^2^). Data represents mean±SD of 3 independent experiments with 6-9 data points ±SD.

Addition of Ag to the sparse nano-ZnO surfaces had transient negative effect on *E. coli* biofilm formation in oligotrophic conditions with an additional 0.5-1.6 log reduction in harvested viable cells (3-48 h post-inoculation, respectively) compared with sparse nano-ZnO without added Ag (Fig. 1a). This additional reduction decreased to a non-significant 0.34 log by 72 h. Ag had only a small but statistically significant effect on *S. aureus* biofilm formation in oligotrophic conditions (<0.5 log additional reduction compared to sparse nano-ZnO) indicating better tolerance to silver compared to *E. coli* (Fig. 1b). As expected, addition of Ag to nano-ZnO surfaces had no effect on bacterial biofilm formation in organic-rich growth medium (Fig. 1d, 1e) due to lower bioavailability of silver.

Nano-ZnO enabled surfaces promoted *C. albicans* biofilm formation in oligotrophic conditions with up to 1.27 log increase in viable attached cells at 48 h time point compared to uncoated surface (Fig. 1c). Silver-containing surfaces had no significant effect on *C. albicans* biofilm formation nor on planktonic viability in oligotrophic conditions compared to uncoated surfaces. However, considering enhanced *C. albicans* biofilm formation on sparse nano-ZnO surfaces and biofilm formation on sparse nano-ZnO/Ag surfaces in oligotrophic conditions demonstrating a significant 0.55-0.72 log reduction compared to sparse nano-ZnO surface during 3-48 h, respectively (Fig. 1c), it could be concluded that silver transiently inhibited *C. albicans* biofilm formation bringing it down to control level. This inhibition was lost 72 h post-inoculation. No biofilm promoting effect was observed in growth medium where inversely a moderate transient biofilm inhibition on both sparse nano-ZnO/Ag (0.85 log decrease) and dense nano-ZnO (0.89 log decrease) surfaces was observed at 48 h (Fig. 1f). Inhibitory effect of silver was further illustrated by the presence of areas with several flattened “ghost cells” on sparse nano-ZnO/Ag surface in growth medium (Supplementary Fig. 1) accompanying slight decrease in viable biofilm cell count 48 h post-inoculation (Fig. 1f). Similar areas with dead cells were not observed in other conditions and seemed to serve as a matrix for the biofilm to grow on.

### Viability of planktonic microbes above the surfaces coated with nano-ZnO or nano-ZnO/Ag

In organics-rich growth media most of the Zn deposited on the nano-ZnO coated surfaces (Fig. 3, a-c) was released into the medium already after 24 h whereas in oligotrophic conditions up to 10 times less Zn was released (Fig. 3, d-f). Increased Zn release in growth media can be attributed to both, culture acidification and formation of protein complexes driving Zn dissolution from nano-ZnO [23], [57], [58]. Higher release of Zn into bacterial growth medium yielded in less biofilm inhibition (Fig. 1d, 1e) and higher viability of planktonic bacteria (Fig. 2d, 2e) while in oligotrophic conditions more nano-ZnO remained on the surface resulting in both more effective inhibition of biofilm formation (Fig. 1a, 1b) and lower planktonic viability (Fig. 2a, 2b). This can possibly be explained by differences in bioavailability. No statistically significant reduction in viability of planktonic yeast cells was registered, regardless of surface type or test media used (Fig. 2c, 2f).

**Figure 2.**
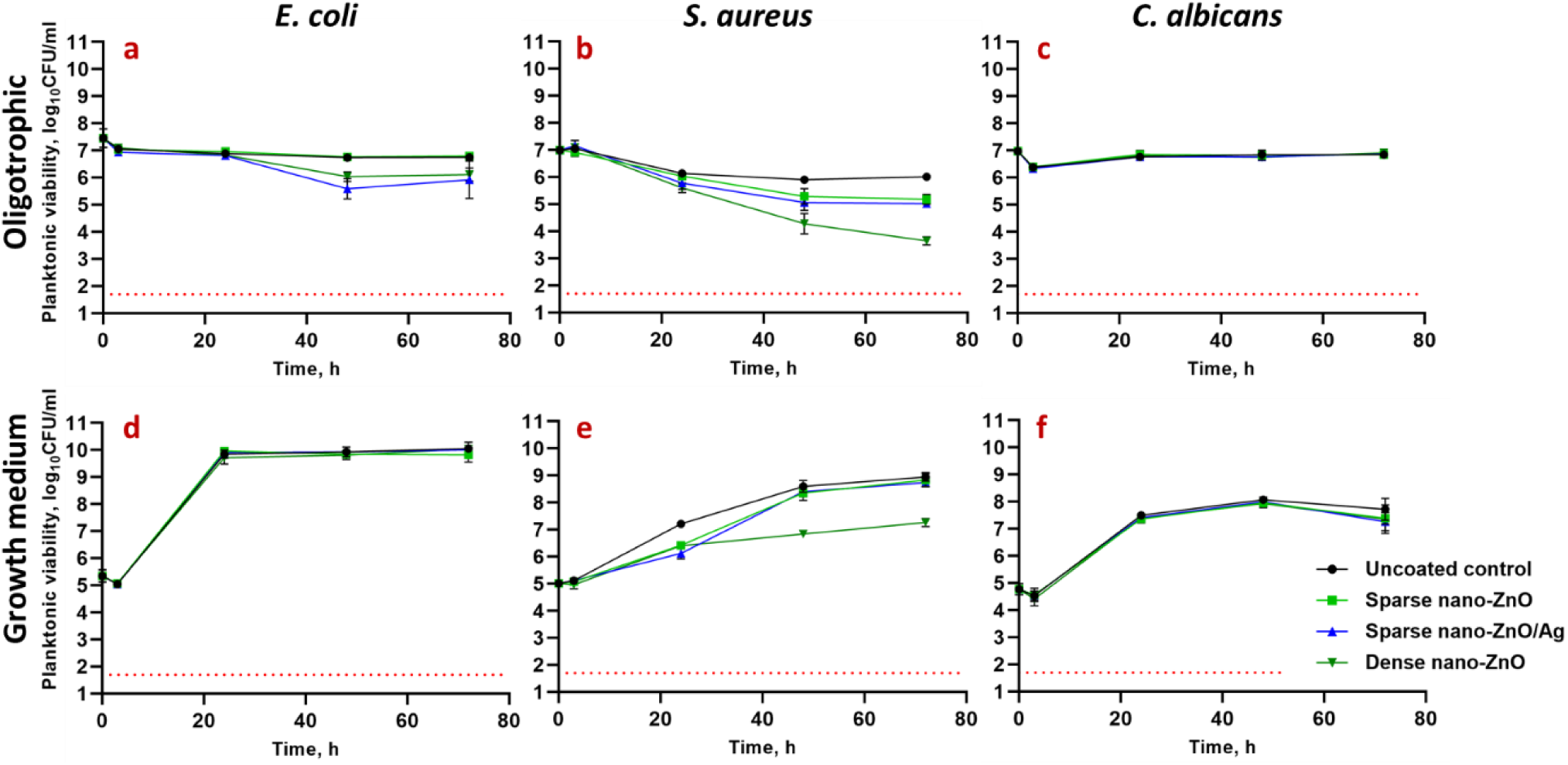
Viability of *E. coli* MG1655 (a, d), *S. aureus* ATCC25923 (b, e) or *C. albicans* CAI4 (c, f) planktonic cells. above nano-ZnO, nano-ZnO/Ag or uncoated surfaces in static oligotrophic environment (1:500 diluted nutrient broth in synthetic tap water; a, b, c) or growth medium (LB: d, e; YPD: f). Planktonic cells were generally more affected by nano-enabled coatings in oligotrophic environment than in growth medium. Red dotted line represents the limit of quantification (50 CFU/ml). Data represents mean±SD of 3 independent experiments with 6-9 data points ±SD.

**Figure 3.**
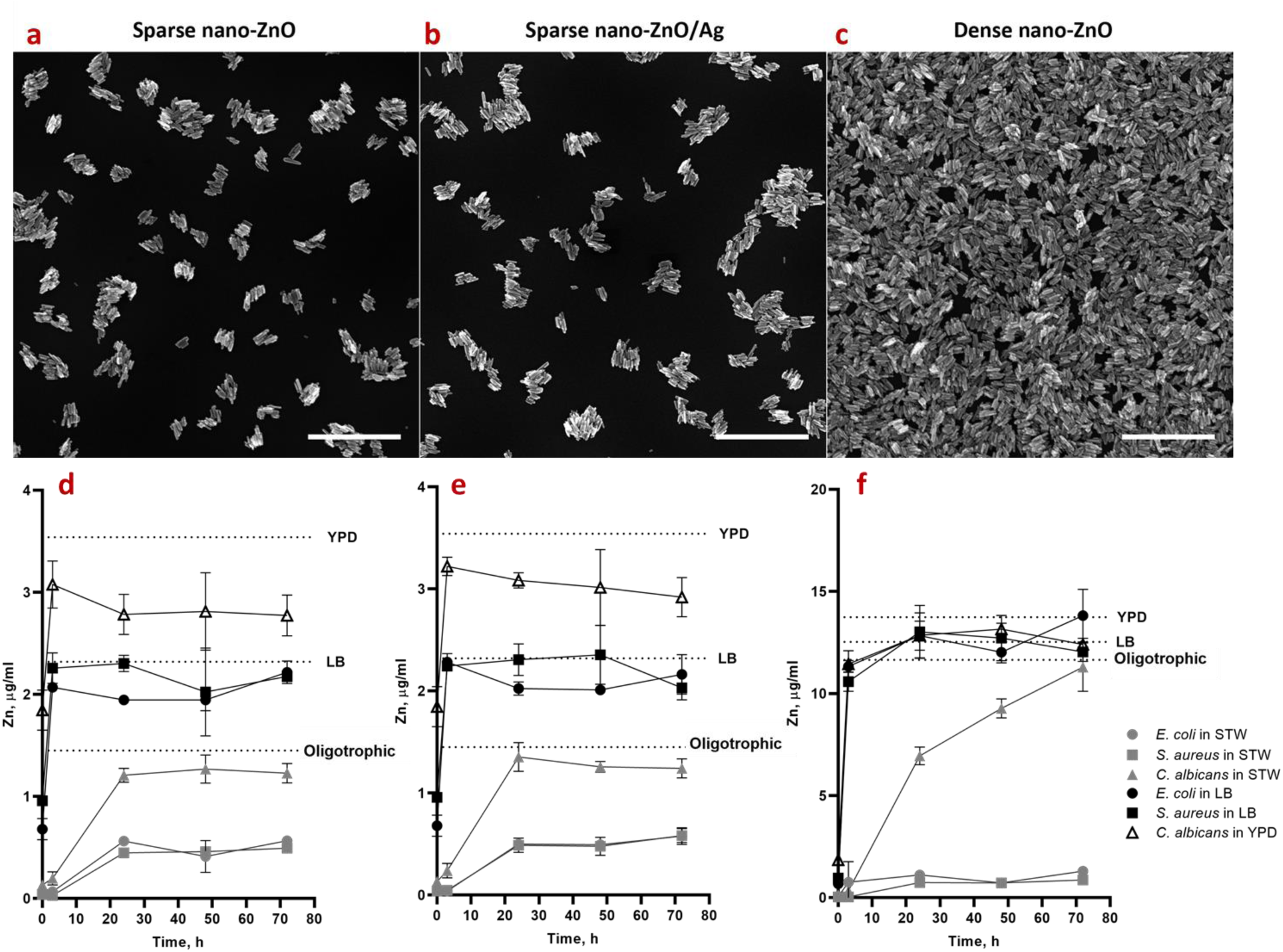
SEM images of sparse nano-ZnO (a), sparse nano-ZnO/Ag (b) and dense nano-ZnO (c) coated surfaces as well as measured Zn release from sparse ZnO (d), sparse ZnO/Ag (e) and dense ZnO (f) surface to the test environments (including the microbes) over time. a, b, c: scale bars represent 1 µm. d, e, f: maximum Zn concentration in each test environment (measured Zn background in different media + theoretical total Zn release from ZnO nanoparticles) is marked as a dotted line. Almost total Zn dissolution was reached already at 3 h in growth medium while dissolution equilibrium was much more slowly achieved in oligotrophic conditions and at lower Zn concentration (d-f). The only exception was *C. albicans* that significantly enhanced Zn release to near total dissolution in oligotrophic conditions. Data represents mean±SD of 3 independent experiments. Total Ag content of 23.56±3.42 ng/cm^2^ in the sparse ZnO/Ag surface was measured which could result in up to 15.27±2.21 ng/ml concentrations in the 5 ml test volume (Supplementary Table 1).

Slight decrease in overall planktonic viable counts at 3 h (Fig. 2, a-f) can be explained by microbial attachment to solid surfaces, including on the polystyrene well surface which decrease planktonic counts in oligotrophic conditions and is not yet compensated by proliferation during 3 h in growth medium at room temperature.

Zinc release from the dense nano-ZnO surface (Fig. 3c, 3f) had a small but statistically significant negative effect of 0.63 logs reduction by 72 h on planktonic *E. coli* viability in oligotrophic conditions and no effect in growth medium (Fig. 2a, 2d) while the same surfaces decreased *S. aureus* planktonic viability in a dose-dependent manner in both oligotrophic conditions and growth medium (Fig. 2b, 2e) with maximum of 2.37 and 1.69 logs reduction, respectively. This effect can likely be attributed to relative sensitivity of *S. aureus* towards Zn.

Ag from the nano-ZnO/Ag surfaces decreased planktonic viability of *E. coli*, but not *S. aureus* and *C. albicans* in oligotrophic conditions (Fig. 2, a-c). Ag from sparse nano-ZnO/Ag had no significant effect on planktonic viability in growth media (Fig. 2, d-f) which can be explained by lower bioavailability of released Ag^+^ ions due to complexing proteins and chloride in the medium [59]–[61]. Maximum calculated Ag concentration in the test system in case of complete release of Ag from the surface coating was estimated to reach 15.27±2.21 ng/ml (Supplementary Table 1). Actual release could not be reliably measured possibly due to high adsorption of Ag to organic matter in the test medium and/or polystyrene well walls.

### Biofilm and cell morphology

Coverage of the surfaces by 48 h biofilm biomass in epifluorescence microscopy (Fig. 4) and higher magnification SEM images of representative microcolonies (Fig. 5) confirmed morphological and structural differences between biofilms grown on uncoated surfaces in growth medium and oligotrophic conditions as well as between biofilms on nano-ZnO coated and uncoated surfaces.

**Figure 4.**
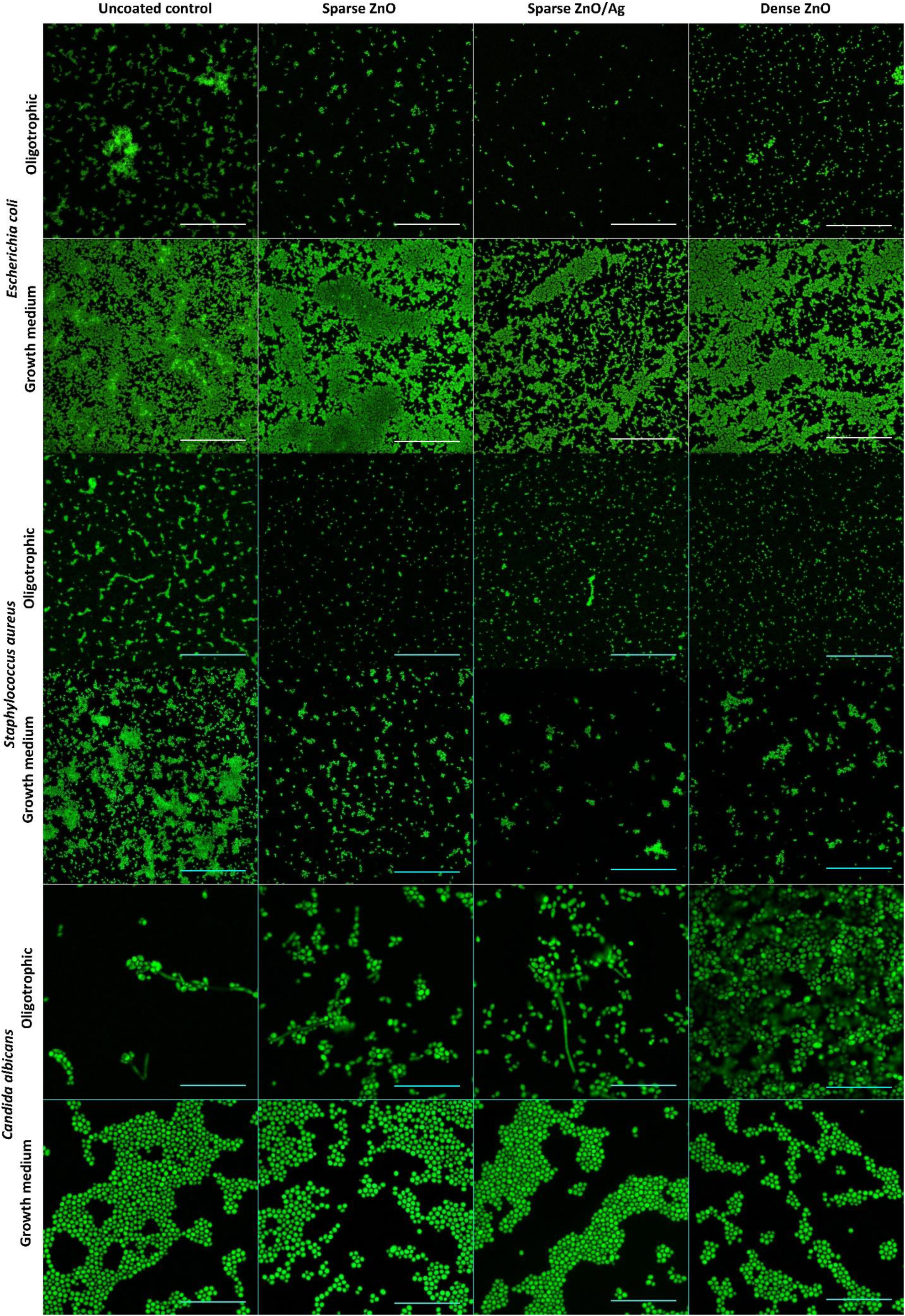
Representative epifluorescence images of *E. coli, S. aureus* or *C. albicans* 48 h biofilms on uncoated surfaces or on surfaces coated with nano-ZnO or nano-ZnO/Ag in oligotrophic conditions and growth medium. Fixed biofilms were stained with DNA- and RNA-binding Syto 9. Scale bars represent 50 µm.

**Figure 5.**
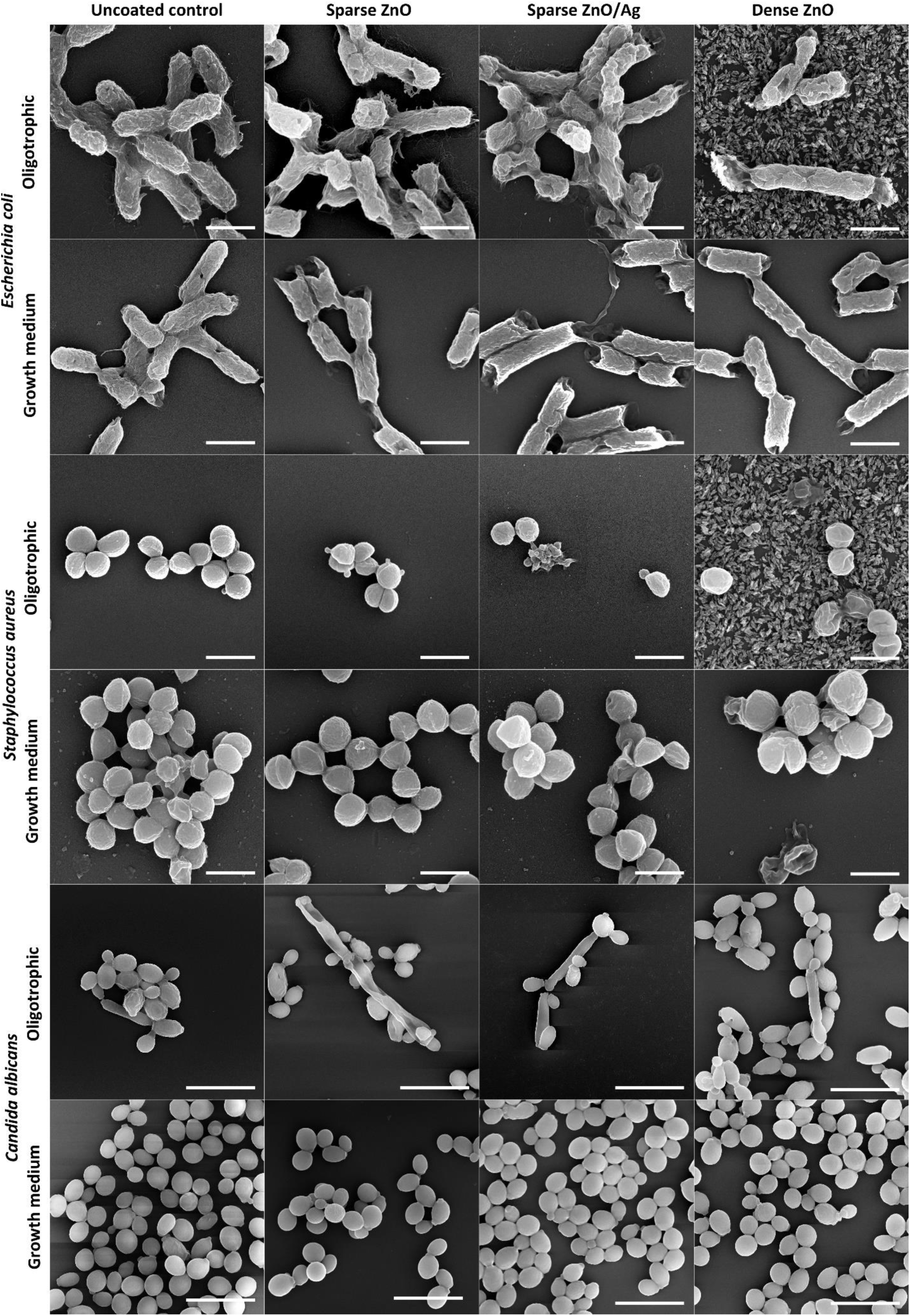
Characteristic SEM images of *E. coli, S. aureus* and *C. albicans* cell and microcolony morphology in 48 h biofilm aggregates on uncoated surfaces or on surfaces coated with nano-ZnO or nano-ZnO/Ag in oligotrophic conditions and growth medium. Scale bars represent 1 µm for bacteria and 10 µm for *C. albicans*. For designation of the surface coatings, see Fig. 3 (a-c).

In case of *E. coli*, also clear differences in cell surface structure were observed. *E. coli* biofilms in growth medium consisted of cells with smoother surfaces and less extracellular matrix (ECM) extended to solid surface while *E. coli* cells in oligotrophic biofilms had a more coarse surface structure and fibrillar ECM extended to solid surface (Fig. 5 upper panels). These differences occurred despite the fact that surface-associated amyloid fibers (SAFs) in the ECM were found to interconnect individual cells of *E. coli* (Supplementary Fig. 2-3) and *S. aureus* (Supplementary Fig. 4-5) biofilms in all growth and exposure conditions used. Although there was less *E. coli* biofilm on nano-ZnO-coated surfaces in oligotrophic conditions compared to uncoated surface (Fig. 1a, Fig. 4 upper panels) and cells tended to be shorter in length on nano-ZnO-coated surfaces, cells with normal morphology could be found even in direct contact with ZnO nanoparticles on dense nano-ZnO surfaces (Fig. 5, upper panels). ZnO nanoparticles were dissolved from the nano-ZnO surfaces in growth medium during 48 h incubation, as also confirmed by elemental analysis (Fig. 3f) and were not visible on SEM images (Fig. 5). Adherence of *S. aureus* cells to nanoparticles on dense nano-ZnO surfaces was also observed, although there were more damaged cells and blebbing, generally associated with cellular damage and virulence [62]–[64], occurring on nano-ZnO surfaces in oligotrophic conditions and on dense ZnO surface in growth medium compared to uncoated surfaces (Fig. 5, middle panels).

There were no clear morphological differences between *S. aureus* biofilm cells on uncoated surfaces in growth medium and oligotrophic conditions besides larger cell size in growth medium (Fig. 5, middle panels). Although Zn-enhanced cell-to-cell adherence has been described for *S. aureus* [65], [66], *S. aureus* tended to preferably form cell aggregates (cell-to-cell adherence) regardless of surface type while *E. coli* tended to cover surface (cell-to-surface adherence) and thereafter form biofilm in height.

Pseudohyphal morphology of the dimorphic fungus *C. albicans* was only observed in oligotrophic conditions, where yeast form was qualitatively more dominant on dense nano-ZnO surface (bottom panels in Fig. 4 and 5). No true hyphae were observed nor expected from this strain in the absence of serum.

### *In vitro* biocompatibility of nano-ZnO and nano-ZnO/Ag surfaces

In order to assess the biocompatibility of nano-ZnO and nano-ZnO/Ag surfaces with human cells, the growth of human keratinocytes directly on the surfaces in standard cell culture conditions was evaluated. Despite the fact that there was enough Zn deposited on dense nano-ZnO surfaces to cause toxicity to the cells in case of total release, amount of Zn released into cell culture medium did not reach cytotoxic concentrations during 48 h (Fig. 6a, 6b) and normal growth and morphology of human keratinocytes on nano-ZnO coated surfaces was observed (Fig. 7). Total Zn and Ag amount deposited on sparsely coated surfaces was not high enough to cause cytotoxicity even in case of theoretical total release into the medium (Fig. 6a, 6b). Compared to control surface, slightly enhanced attachment and growth of keratinocytes on dense nano-ZnO surfaces (Fig. 6c) was observed. This observation cannot be explained by differences in surface properties of the different surfaces, e.g., hydrophilicity (see water contact angles on different surfaces in Supplementary Fig. 6).

**Figure 7.**
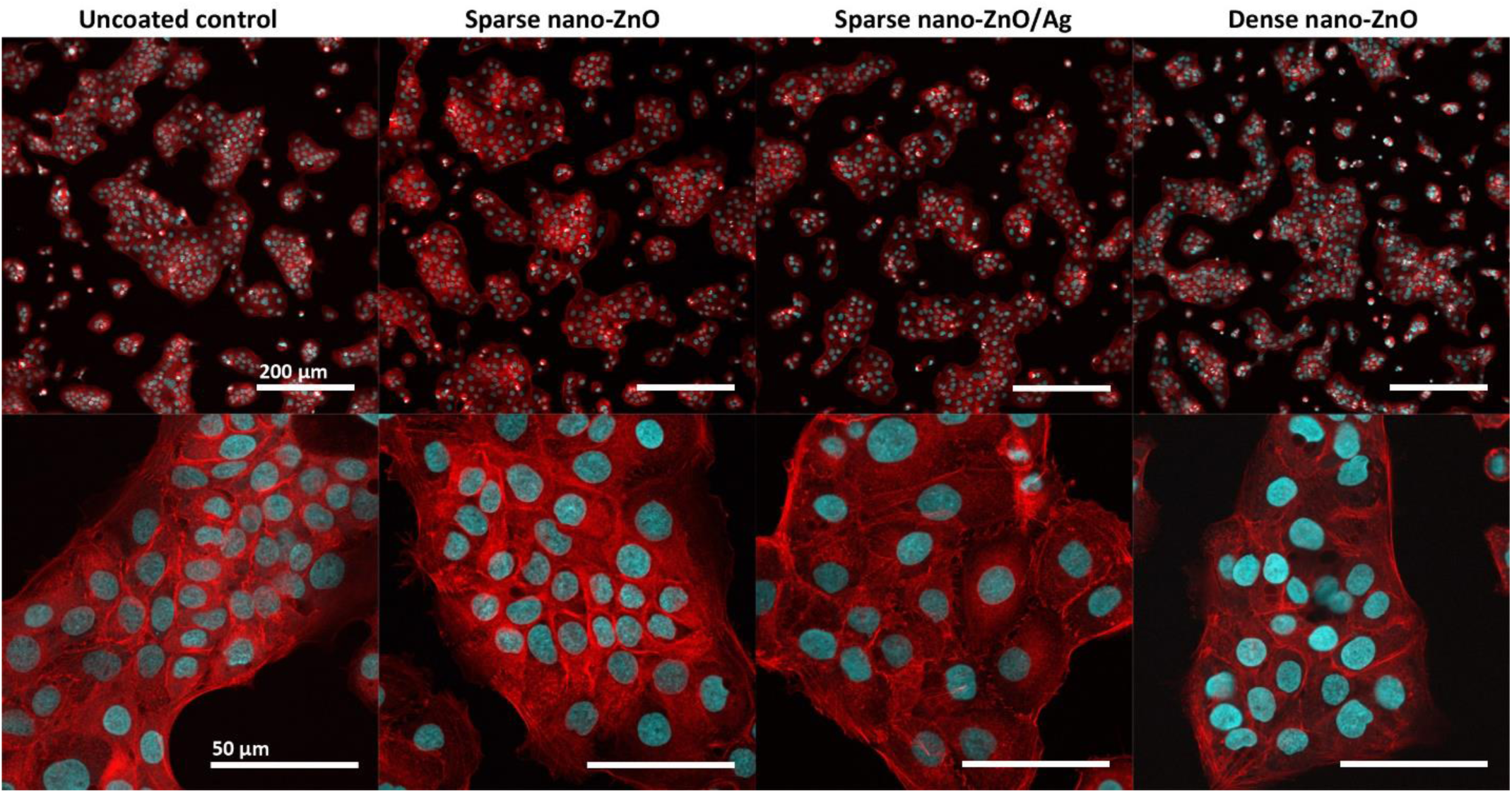
Representative confocal microscopy (CLSM) images of HaCaT keratinocytes on nano-ZnO or nano-ZnO/Ag coated surfaces: upper panel – lower magnification CLSM (scale bar 200 µm), bottom panel – higher magnification CLSM (scale bar 50 µm); nuclei – cyan (Hoechst 33342), cytoskeleton – red (Phalloidin-TRITC). Cells are viable (Fig. 6) and appear normal on all surfaces.

**Figure 6.**
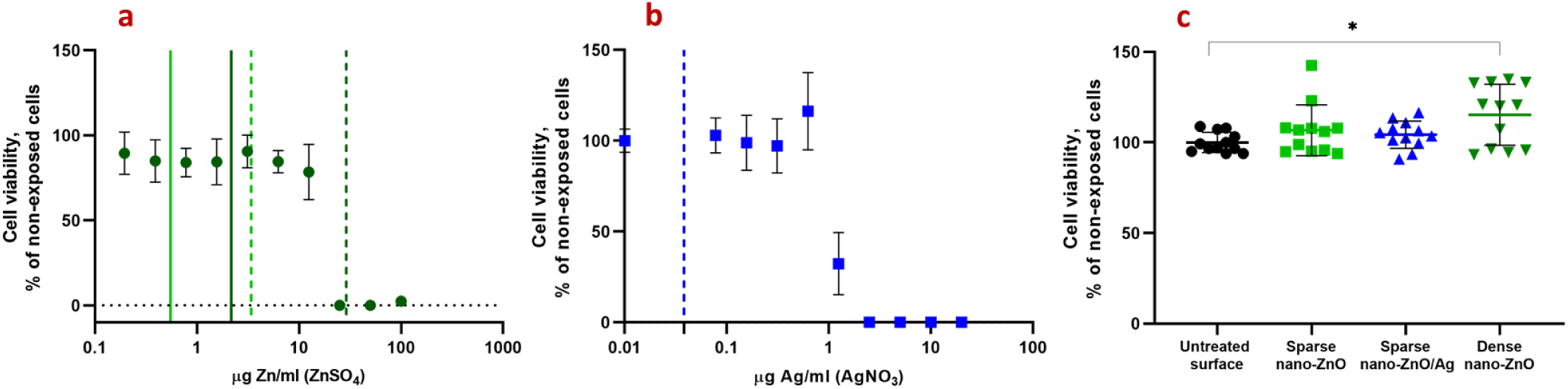
Cytotoxicity of soluble Zn (a) and Ag (b) salts or nano-ZnO and nano-ZnO/Ag coated surfaces (c) to HaCaT keratinocytes after 48 h in standard cell-culture conditions (DMEM high glucose medium supplemented with Na-pyruvate (1mM), penicillin-streptomycin (100 U/mL;100 μg/mL), fetal bovine serum (10%), 37 °C, 5% CO_2_). **a:** Viability of HaCaT cells exposed to ZnSO_4._ Solid line denotes Zn released into cell culture medium from sparse nano-ZnO (light green) and dense nano-ZnO (dark green) surfaces during 48 h. Dotted lines denote respective maximal theoretical concentration in case of total release of deposited Zn. **b:** Viability of HaCaT cells exposed to AgNO_3._ Dotted blue line marks theoretical maximum release of Ag into the test environment from sparse nano-ZnO/Ag surface. Actual release could not be reliably measured possibly due to high adsorption of Ag to organic matter in the test medium and/or polystyrene well walls. **c:** Cytotoxicity of nano-ZnO and nano-ZnO/Ag coated surfaces to HaCaT keratinocytes cultured directly on the surfaces. No direct cytotoxicity of nano-enabled surfaces was observed. Increased growth of keratinocytes was observed on dense nano-ZnO surfaces. * denotes statistically significant difference (P<0.05). Zn release from the surfaces during 48 h did not reach cytotoxic range. Theoretical maximal Zn concentration resulting from total Zn release from the dense nano-ZnO surfaces was in toxic range while total Zn from sparse nano-ZnO and Ag from sparse nano-ZnO/Ag would not have reached toxic concentrations in cell culture medium.

## DISCUSSION

The extracellular matrix (ECM) is an integral part of biofilms and can make up most of the volume of a mature biofilm [68] but total biomass or biovolume is not a good proxy for total or viable cell count in a biofilm, especially when comparing chemical treatments or carrier surfaces that could potentially affect ECM composition. However, viable microbes are critically needed to establish biofilms *de novo* and the sole cause of pathogen transfer via high-touch surfaces. Therefore, we chose to quantify viable microbes harvested from 3-72 h biofilms to evaluate *de novo* surface colonization. As harvested CFU counts exclude ECM quantification entirely, we also evaluated the biofilms qualitatively by epifluorescence and electron microscopy. Epifluorescence microscopy was chosen to be performed on 48 h fixed biofilm samples monostained with Syto 9 instead of viable biofilms stained with Syto 9 and propidium iodide (PI) because (i) based on pilot experiments, largest effect was expected at 48 h; (ii) PI is known to underestimate viability in biofilms [9], [10]; (iii) highly variable Syto 9 intensity in viable biofilms depending on viability state and Gram type of the cells [9], [69]; (iv) time limits of working with viable samples to achieve comparable representative results.

It must be noted, however, that adding liquid manipulation steps such as rinsing, fixation and gradual dehydration to the preparation of biofilms is likely to affect the appearance of biofilms and cause e.g., loss of biomass in each step. Kragh *et al.* have recently demonstrated that media removal and rinsing can critically affect quantitative biofilm analysis compared to undisturbed biofilms [10]. In addition, we observed that while oligotrophic bacterial biofilms retained their overall microcolony architecture during fixing and dehydration steps, biofilms formed in growth medium lost most of their biomass and surface coverage during dehydration steps prior to SEM. Taking into account also the rapid biofilm formation during the first 3 h in oligotrophic conditions (Fig. 1) we empirically suggest that biofilms formed in oligotrophic conditions are more strongly attached to the substratum.

Reductions in harvested viable counts of bacterial biofilms on nano-ZnO and nano-ZnO/Ag surfaces were accompanied by respective reductions in planktonic viable counts regardless of media used and can therefore be mostly attributed to general antibacterial activity of the Zn-enabled surfaces and metal dissolution in concordance with previous knowledge [13], [70] and not biofilm-specific mode of action. *S. aureus* was more sensitive to Zn and *E. coli* more sensitive to Ag for both biofilms and planktonic cultures in accordance with our previous results using a planktonic test in oligotrophic conditions [53]. *S. aureus* cells attached to ZnO-coated surfaces (Fig. 5, middle panels) appeared damaged on SEM micrographs and presented blebs. Blebbing has been described not only for Gram-negative, but also for Gram-positive bacteria and fungi [62] and appears to have a more general role in stress response, virulence and intercellular communication across the domains of life. Vesicle formation has also been confirmed in case of *S. aureus* [63] and can take place for example in response to antibiotic stress thereby increasing virulence [64].

We have shown previously that choice of media critically affects metal toxicity [61] and indeed, it was evident, that both Zn and Ag were more toxic in oligotrophic conditions compared to no effect on either biofilm or planktonic cells of *E. coli* in growth medium. This can be explained by lower metal bioavailability in organics-rich growth medium [57]–[61] but one must acknowledge that also physiology of exponentially growing and starving non-dividing microbes is different. Although classical microbiological culture media are the most used testing environments, their suitability for each application should be evaluated.

Although metal effect to surface associated amyloids (SAFs) in biofilm matrix is yet scarcely studied [44], [45], we also hypothesized that high Zn concentrations might interfere with bacterial surface-associated amyloid [71]–[74] formation as it can negatively affect fibrillar structure of other amyloids [75]–[79] and could thereby disrupt SAF-mediated surface colonization. Although cell surface morphology seemed to differ between *E. coli* cells grown on nano-ZnO and uncoated surfaces in oligotrophic conditions (Fig. 5, top panel), we found no substantial qualitative differences in amyloid staining of those biofilms (Supplementary Fig. 2-5). Qualitative results do not strictly rule out smaller Zn-dependent differences, which amyloid signal quantification could possibly discriminate, if present.

While reduction in bacterial biofilm viability was accompanied by respective antibacterial activity against planktonic cells there were clear biofilm-specific effects on *C. albicans* biofilm formation. Although planktonic viable counts were not significantly different from uncoated control surface regardless of surface type or media used, biofilm formation was affected. Enhanced dose-dependent biofilm formation of *C. albicans* on nano-ZnO-coated surfaces in oligotrophic conditions cannot be explained by differences in the hydrophilic properties of nano-ZnO coated and uncoated surfaces (Supplementary Fig. 6). However, *C. albicans* has been demonstrated to possess systems to harvest extracellular Zn and store Zn intracellularly in zincosomes [80] which enables *C. albicans* to both overcome nutritional immunity in host organism as well as to detoxify Zn^2+^ excess. Active uptake of extracellular Zn could also be the reason, why nano-ZnO was depleted from the surfaces in oligotrophic conditions in the presence of *C. albicans* but not bacteria (Fig. 3, d-f). For dense nano-ZnO surfaces this effect was also clearly observable on SEM images (Fig. 5, right column). Interestingly, previously published results on zinc effects on *Candida* biofilm formation are contradictory. For example, Kurakado *et al.* demonstrated an increase in *C. albicans* biofilm formation as a response to ZnSO_4_ and a decrease in response to zinc chelator [81] while several others report Zn-associated biofilm inhibition [82]–[84]. These contradictions could be partly due to different average and local Zn concentrations reached in these studies while we had the nano-ZnO attached to a solid surface resulting in lower average Zn concentration in the test environment or due to growth-related and physiological differences in various oligotrophic and nutrient-rich environments as well as differences in yeast strains used. We also observed that pseudohyphal morphology of *C. albicans* [67] was only registered in oligotrophic conditions where yeast form was qualitatively more dominant on dense nano-ZnO surface (bottom panels in Fig. 4 and 5). Harrison *et. al* have demonstrated that sub-inhibitory Zn^2+^ concentrations can indeed interrupt yeast to hyphal differentiation in *Candida* biofilms [85] and it has been suggested that zinc-responsive transcription factor Zap1 is a negative regulator of biofilm matrix accumulation and maturation [86] also directing the yeast-hypha balance to yeast form [87]. On the other hand, filamentation is not a prerequisite for initial biofilm formation [88] as can also be seen in our results.

*Candida* spp is an important cause of nosocomial bloodstream infections [90]. Although we did not look into polymicrobial biofilms, enhancement of *C. albicans* biofilm formation could be especially disadvantageous in biomedical applications as many patients with candidemia tend to present polymicrobial blood cultures [91] and surface colonization by *C. albicans* can in turn enhance polymicrobial biofilm formation or drug tolerance of other potential pathogens [92]–[94]. In the light of existing and emergent drug-resistant *Candida* spp. [89] Zn-enhanced *C. albicans* biofilm formation in oligotrophic conditions should be studied further to reveal mechanistic causes of the biofilm-promoting phenomenon.

Zinc oxide is well tolerated by humans and is listed as a generally recognized as safe (GRAS) substance by the US Food and Drug Administration (21CFR182.8991), approved for use in cosmetics in EU for up to 25 % concentration in the ready-for-use preparation (Regulation (EC) No 1223/2009) and widely used in topical applications (incl. sunscreens, baby powders, diaper creams, mineral make-up etc.) as well as food additive, corrosion protection, filler/pigment in paints alongside titanium dioxide etc. We would like to emphasize that growing human keratinocytes directly on the surfaces of interest as a measure of biocompatibility was used as a worst-case-scenario during material development phase. *In vivo* dermal exposure to surface-bound ZnO is expected to be lower due to the lack of fully functional skin barrier in cell culture conditions and shorter exposures by high-touch surfaces. The observed slightly higher viability of human keratinocytes on dense nano-ZnO surfaces is not entirely unexpected as zinc is known to promote wound healing [95] and has been shown to also promote keratinocyte migration and proliferation as well as to modulate integrin expression needed for adherence to extracellular matrix [96]. However, the lack of any cytotoxicity for keratinocytes on ZnO and ZnO/Ag surfaces does not rule out potential sublethal events, e.g., genotoxicity that has been observed at somewhat lower zinc concentrations than cytotoxicity [97], [98] and these specific toxic effects would require further studies.

## CONCLUSION

This study demonstrated that in general, bacterial *E. coli* and *S. aureus* monospecies biofilm formation on nano-ZnO and nano-ZnO/Ag composite-enabled surfaces was inhibited in a dose-dependent manner: sparsely coated nano-ZnO surfaces were less inhibitory than densely coated ones and inhibition of biofilm formation was accompanied by antibacterial activity against planktonic cells. The surfaces were not cytotoxic to human keratinocytes and no significant inhibition of yeast biofilm was observed. On the contrary, enhancement of *C. albicans* biofilm formation was registered on densely coated nano-ZnO surfaces in oligotrophic conditions. This observation is certainly disadvantageous considering the emergence of drug-resistant infections of *Candida spp.* and yeast role in polymicrobial biofilms. Our results indicate that best approach to achieve broad-spectrum inhibition of biofilm formation on high-touch surfaces would be the combination of ZnO and silver. It is also important to highlight the role of the test environment. Except for Zn toxicity towards *S. aureus*, all above-mentioned antimicrobial and antibiofilm effects were detected only in oligotrophic environment and not in widely used microbial growth media. Moreover, selection of detection method proved critical in the quality of information acquired as biomass was lost in every liquid manipulation step. Media removal and washing steps have been shown to critically affect quantitative biofilm analysis. We add to that and propose that liquid manipulation steps can also easily introduce compositional biases e.g. exaggerate biomass differences formed in different conditions.

## METHODS

### Surfaces

The preparation of ZnO and ZnO/Ag nanoparticles is more thoroughly described in our previous work [53]. Briefly, hydrothermal synthesis was used to prepare ZnO nanoparticles using zinc acetate as a precursor. ZnO/Ag composite particles were created by supplementing ZnO nanoparticles with Ag using UVA-induced photodeposition. ZnO or ZnO/Ag nanoparticles were deposited on 18×18 mm square cover glasses (2855-18, Corning) using spin-coating and subsequently annealed to enhance particle surface attachment. Two different coating densities of nano-ZnO were used further referred to as sparse and dense, depicted on Fig. 3 (a-c). Nano-ZnO/Ag composite particles were used only in sparse coating density. Dense coating resulted in uniformly covered surfaces while sparse coatings exposed glass carrier surface with distance between nanoparticle clusters comparable to bacterial cell size as can be seen on SEM images of the surfaces on Fig. 3. Physical and chemical properties of the nanoparticles used as well photocatalytic and antibacterial activity of the surfaces used in this study have been previously described [53]. Total Zn content in the nano-coatings was analyzed using TXRF (S2 Picofox, Bruker) and Ag content with AAS (ContrAA 800, Analytic Jena AG) after 1 h treatment in 0.5 ml concentrated HNO_3_. The results are shown in Supplementary Table 1.

### Biofilm formation, harvesting and quantification

Monospecies biofilms of *E. coli* MG1655, *S. aureus* ATCC25923 and *C. albicans* CAI4 were studied. Sterile nano-ZnO, nano-ZnO/Ag or uncoated surfaces were placed into non tissue culture treated 6-well polystyrene plate wells. 5 ml of microbial inoculum was added to each well. To prepare inoculums, overnight cultures of microbes were pelleted by centrifugation (7000 g, 10 min) and washed twice with deionized water before re-suspension in test medium and density optimization. Initial cell density was about 10^7^ CFU/ml in oligotrophic conditions, i.e., synthetic tap water (STW; Table 1) containing 1:500 diluted nutrient broth (NB; Table 1) and 10^5^ CFU/ml in growth media, i.e., lysogeny broth (LB, Table 1) in case of bacteria and yeast extract-peptone-dextrose medium (YPD, Table 1) in case of yeast. Surfaces were incubated in dark static conditions (covered with aluminum foil) at room temperature (22 °C) to allow biofilm formation. After 0, 3, 24, 48 and 72 h three samples of each surface were dip-rinsed in PBS (Table 1) with sterile forceps, carefully drained from one corner and submerged in 15 ml 0.5x neutralizing broth (SCDLP; Table 1) with 1.5M NaCl in 50 ml centrifuge tube, vortexed for 30 sec at max speed, ultrasonicated for 15 min in ultrasonic bath (Branson1800) and again vortexed for 30 sec. An aliquot of each resulting suspension was serially diluted in PBS and drop-plated on nutrient agar (NA) or yeast growth agar (YPD agar, Table 1) for colony counting. Planktonic fraction in 5 ml media over biofilms in each well was mixed by pipetting, serially diluted and plated for planktonic colony count. To assess Zn release from the surfaces, an aliquot from planktonic fraction (including microbes) in wells above the biofilms was taken for elemental analysis. TXRF (S2 Picofox, Bruker) was used for Zn analysis. Maximum calculated Ag concentration in the test system in case of complete release of Ag from the surface coating was estimated to reach 15.27±2.21 ng/ml (Supplementary Table 1). Actual release could not be measured due to high adsorption of Ag to organic material and well walls.

**Table 1.** Media and buffers.

|  |  |
| --- | --- |
| Synthetic tap water (STW; DIN10531), prepared as 5x stock | 300 mg/L NaHCO<sub>3</sub>, 175 mg/L MgSO<sub>4</sub>, 300 mg/L CaCl<sub>2</sub>; pH 7.5 in 1xSTW |
| Nutrient broth (ISO 22196) | 3 g/L meat extract, 10 g/L peptone, 5 g/L NaCl; pH 6.8-7.2 |
| Lysogeny broth (LB) | 5g/L tryptone, 5 g/L yeast extract, 5 g/L NaCl; pH 6.8-7.2 |
| Yeast extract peptone dextrose medium (YPD) and respective agar | 20 g/L peptone, 10 g/L yeast extract, 20 g/L glucose + 15 g/L agar for plates; pH 6.8-7.2 |
| Phosphate buffered saline (PBS) | 8 g/L NaCl, 0.2 g/L KCl, 0,81 g/L, Na<sub>2</sub>HPO<sub>4</sub>x2H<sub>2</sub>O, 0.2 g/L KH<sub>2</sub>PO<sub>4</sub>; pH 7.1 |
| Nutrient agar (ISO 22196) | 5 g/L meat extract, 10 g/L peptone, 5 g/L NaCl, 15 g/L agar; pH 7.0-7.2 |
| Soybean casein digest broth with lecithin and polyoxyethylene sorbitan monooleate (SCDLP broth; ISO 22196) | 17 g/L casein peptone, 3 g/L soybean peptone, 5 g/L NaCl, 2.5 g/L Na<sub>2</sub>HPO<sub>4</sub>, 2.5 g/L glucose, 1.0 g/L lecithin, 7.0 g/L nonionic surfactant; pH 6.8-7.2 |
| Sodium cacodylate buffer, prepared as 0.2 M stock | 4.8 g Na(CH<sub>3</sub>)<sub>2</sub>AsO<sub>2</sub> x 3 H<sub>2</sub>O in 100 ml deionized water brought to pH 7.4 with the addition of 5.6 ml 0.2M HCl |

Viable counts of cells harvested from biofilms were calculated as follows:

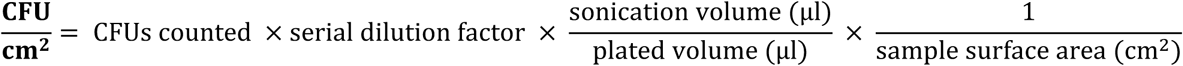

Viable counts of inoculums and planktonic cells were calculated as follows:

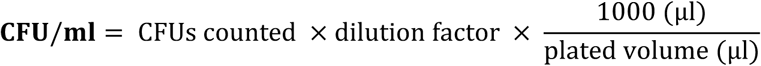

### Ultrasonication protocol

SCDLP (Table 1) was used for toxicity neutralizing properties (ISO 27447) and its surfactant content was expected to enhance releasing biofilms from surfaces. 1.5M NaCl was used to aid in biofilm harvesting as it has also been successfully used for extracellular matrix removal [99] and does not influence microbial viability when combined with ultrasonication and vortexing (Supplementary Fig. 7). Staining with 30 µM propidium iodide and 5 µM Syto 9 with subsequent epifluorescence microscopy (see below) was used to evaluate biofilm harvesting efficiency. It was confirmed that adherent cells were removed from the surfaces (data not shown).

### Staining and microscopy

Biofilms were stained to visualize total cellular biomass (DNA/RNA), surface-associated amyloid fibers (SAF) and chitin (in case of yeast). Before staining, the biofilms were dip-rinsed in PBS (Table 1) to remove loosely bound cells as described above and fixed by 2-h incubation in 2.0% glutaraldehyde in 0.1M sodium cacodylate buffer (Table 1) followed by rinsing once in the same buffer. For total cellular biomass visualization, nucleic acids were stained by incubating the fixed cells for 15 min with 5 µM Syto 9. For staining of SAF and chitin (fungal cell wall component), fixed biofilms were incubated for 15 min with 5 µM Syto 9 and 10 µM Congo Red. Stained biofilms were visualized with Olympus CX41 using exciter filter BP475, dichroic mirror DM500, barrier filter O515IF (Syto9 staining) or with Zeiss LSM800 confocal laser scanning microscope (CLSM) using excitation/emission track settings of 488 nm/505-550 nm (Syto 9) and 561 nm/575-700 nm (Congo Red).

Congo Red (CR), used to stain SAFs, also stains cellulose and chitin in fungal cell walls [100] due to structural similarity to cellulose. In this study, we have used CR also to visualize *C. albicans* cells by staining both cell wall and fungal SAFs [101]. Regarding possible cellulose signal interfering with amyloid staining of *E. coli*, as cellulose is a known component in ECM of several *Enterobacteriaceae* species, it is known that *E. coli* K-12 strains, including MG1655 have a stop codon mutation in the cellulose operon and do not produce cellulose [102].

To visualize biofilm morphology, electron microscopy was used. For that, fixed biofilms were gradually dehydrated with ethanol, air-dried, coated with 3 nm Au/Pd layer (Polaron Emitech SC7640 Sputter Coater) and visualized with scanning electron microscopy (FEI Helios NanoLab 600).

### Cytotoxicity testing

Toxicity of the surfaces was tested on human immortalized HaCaT keratinocytes (ThermoFisher). Cells were maintained in DMEM high glucose medium supplemented with Na-pyruvate (1mM), penicillin-streptomycin (100 U/mL and 100 μg/mL, respectively) and fetal bovine serum (10%) at 37 °C and 5% CO_2_. During maintenance, the cell density was kept between 0.5-1×10^6^ cells/ml. Each type of surface in four replicates was placed into a separate well of a 6-well plate and 2 ml of cell suspension (4*10^4^ cells/mL) in supplemented DMEM medium was added. Cells were cultivated for 48 h, surfaces were removed using tweezers to a new well of a 6-well plate. An aliquot of the remaining medium was taken for Zn concentration measurement by TXRF (S2 Picofox, Bruker). One replicate of each type of surface was left for microscopy while to the rest of three replicates 0.33% of Neutral Red (Sigma) solution in supplemented DMEM medium was added and incubated at 37 °C and 5% CO_2_ for 4 hours. After that, the dye solution was removed by double washing with PBS and the dye remaining within viable cells was dissolved over 10 min (gentle shaking) using 1 ml of Neutral Red Assay Solubilization Solution (Sigma). The intensity of Neutral Red dye was measured spectrophotometrically at 540 nm using Fluoroskan plate reader (Thermo). Cell viability on nano-ZnO and nano-ZnO/Ag surfaces was calculated as percentage of viability on uncoated surface. Cell viability assays on surfaces were repeated twice. For microscopy, the surfaces with cells were first covered with 3.7% formaldehyde in 0.1% TritonX-100 solution for 10 min, then with 1 µg/ml solution of Hoechst 33342 (ThermoFisher) and 1 µg/ml Phalloidin-TRITC (Sigma) in PBS for 30 min. After that, the dye solution was removed and cells were imaged using confocal microscope with filter settings for DAPI and TRITC.

For cytotoxicity assays of respective soluble salts (Fig. 7), exposure to HaCaT cells was carried out in 96-well format and cell viability was assessed using Neutral Red assay, essentially as described above.

### Statistical analysis

One- or two-way ANOVA analysis followed by Tukey *post-hoc* test was used to detect significant differences in multiple comparisons at 0.05 significance level using GraphPad Prism 8.3.0. Log-transformed data was used for the analysis of CFU counts. Only statistically significant (P<0.05) differences and log reductions are mentioned in the text unless stated otherwise.

## Funding & acknowledgements

We thank Arvo Tõnisoo for help with SEM imaging and Dr Nuno F. Azevedo for consulting on biofilms. This work was supported by the Estonian Research Council grants IUT 23-5, PUT 748, EAG20, PRG749, European Regional Development Fund projects “Emerging orders in quantum and nanomaterials (TK134)” and “Advanced materials and high-technology devices for sustain-able energetics, sensorics and nanoelectronics (TK141)” and ERDF project Centre of Technologies and Investigations of Nanomaterials (NAMUR+, project number 2014-2020.4.01.16-0123).

## Contributions

MR designed and executed biofilm experiments, analyzed the data and wrote the manuscript. MV designed and prepared the surfaces. AI designed and executed the cell culture experiments, assisted in experimental design, participated in writing and amending the manuscript. HV consulted on and executed elemental analysis. KK consulted regarding yeast biology and assisted in writing the manuscript. VK designed the surfaces, assisted in experimental design, participated in writing and amending the manuscript. AK consulted on general microbiology and critically edited the manuscript.

## Supplementary figures

**Supplementary Figure 1.**
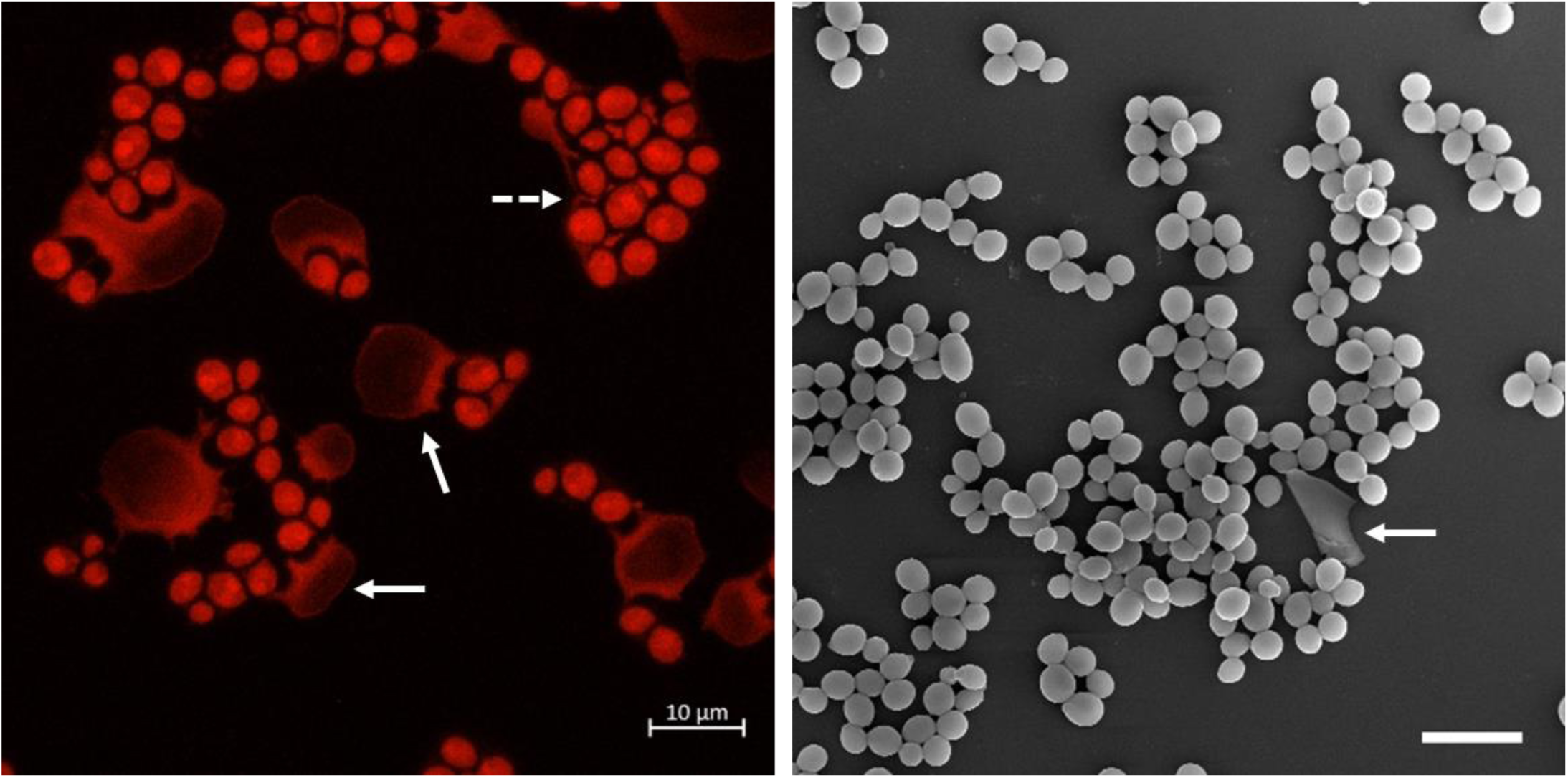
48 h *C. albicans* biofilm on sparse ZnO/Ag surface in growth medium: CLSM maximal orthogonal projection with cell walls stained red with Congo red (left) and SEM image (right). White arrows indicate flattened dead “ghost cells” that occur in patches on nano-ZnO/Ag surfaces and seem to act as a carrier for biofilm cells (dotted arrow). Most “ghost cells” are laterally detached from neighboring cells during several liquid manipulation steps in the fixing protocol and are mostly lost in SEM images. Scale bars represent 10 µm.

**Supplementary Figure 2.**
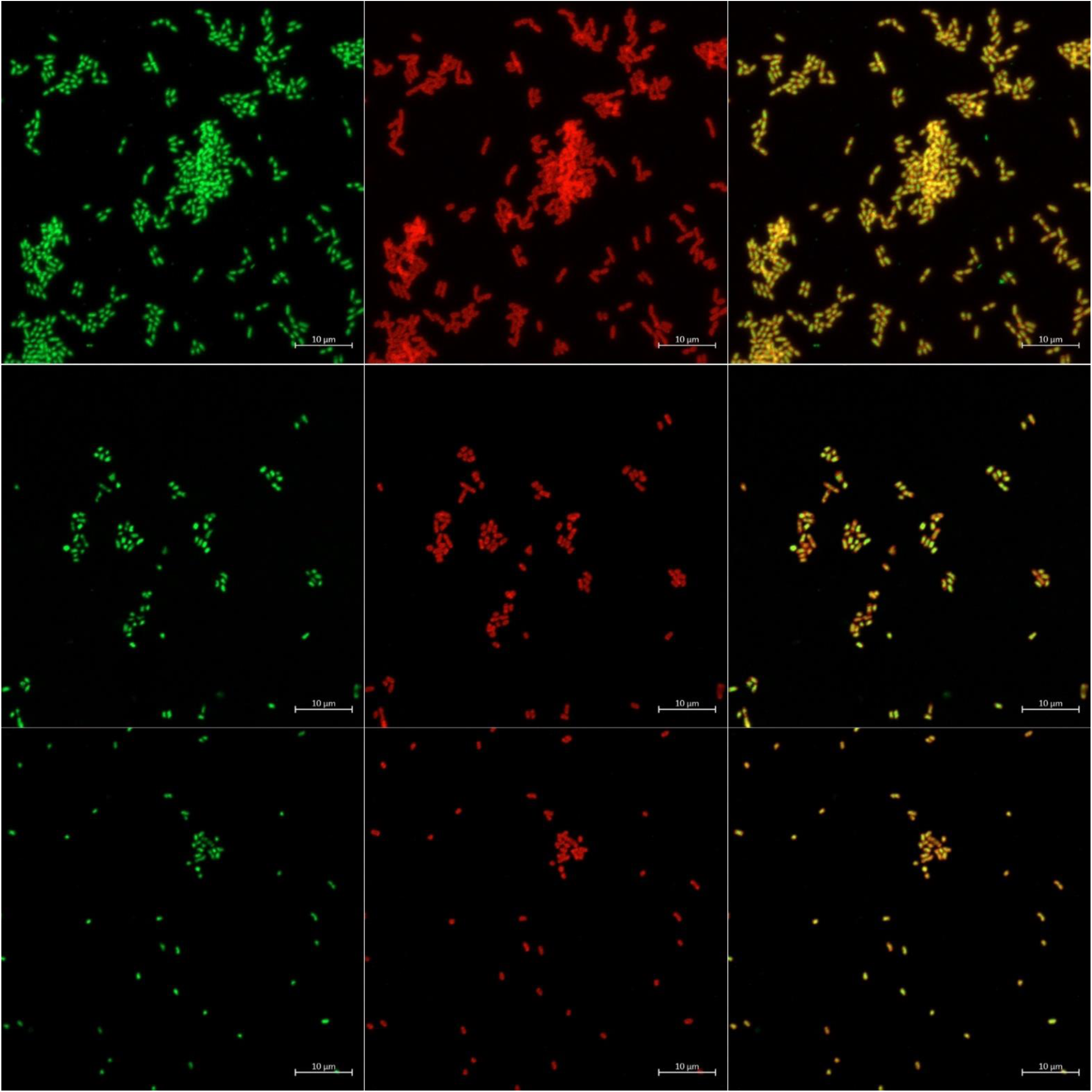
Maximal orthogonal projections of representative CLSM images of fixed 48 h oligotrophic *E. coli* biofilm on uncoated surface (upper panel), sparse nano-ZnO surface (middle panel) and dense nano-ZnO (bottom panel). DNA/RNA stained with Syto9 (green channel, left column) and surface-associated amyloid fibers stained with Congo Red (red channel, middle column). Combined channel view in right column. Scale bars represent 10 µm.

**Supplementary Figure 3.**
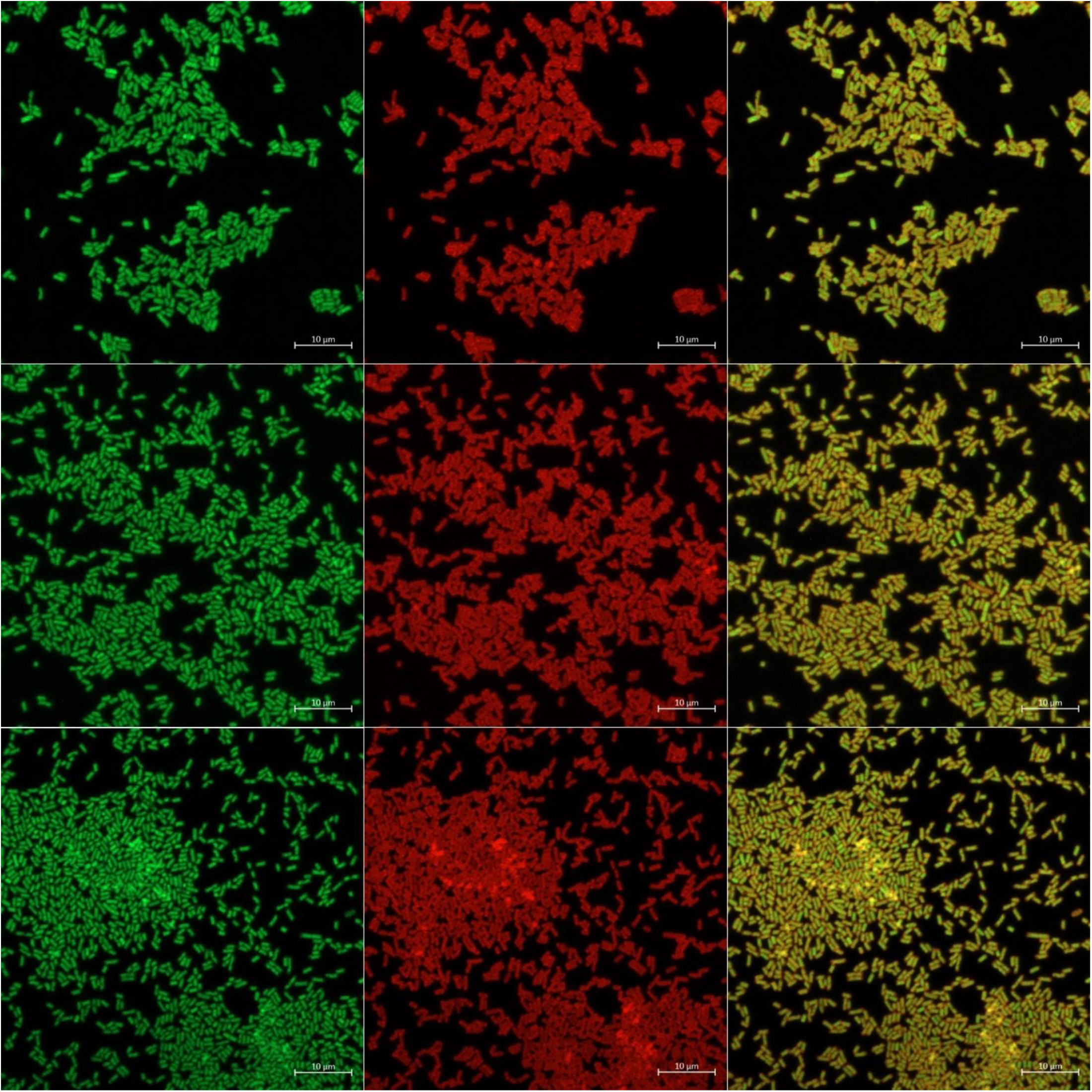
Maximal orthogonal projections of representative CLSM images of fixed 48 h growth medium *E. coli* biofilm on uncoated surface (upper panel), sparse nano-ZnO surface (middle panel) and dense nano-ZnO (bottom panel). DNA/RNA stained with Syto9 (green channel, left column) and surface-associated amyloid fibers stained with Congo Red (red channel, middle column). Combined channel view in right column. Scale bars represent 10 µm.

**Supplementary Figure 4.**
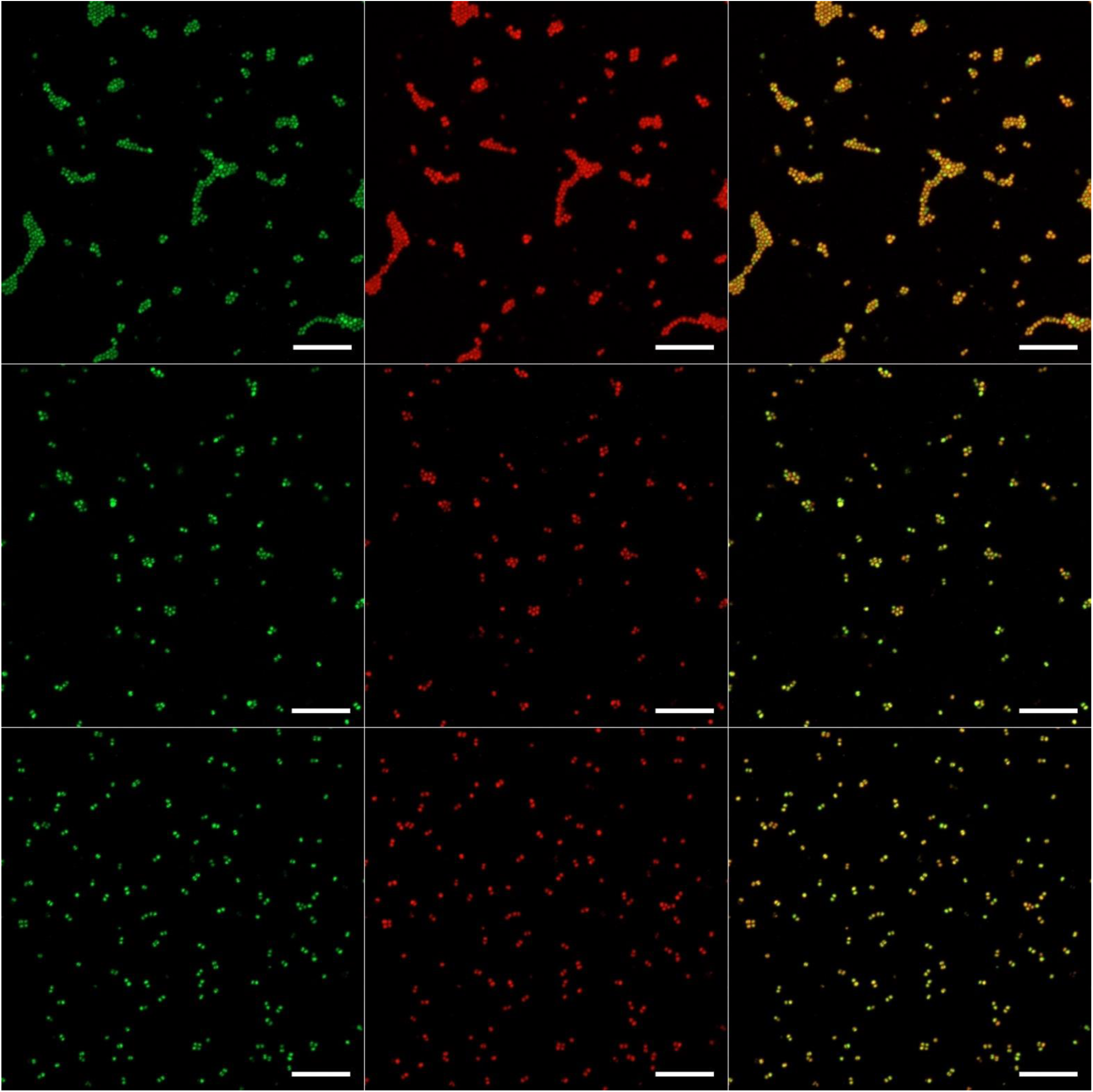
Maximal orthogonal projections of representative CLSM images of fixed 48 h oligotrophic *S. aureus* biofilm on uncoated surface (upper panel), sparse nano-ZnO surface (middle panel) and dense nano-ZnO (bottom panel). DNA/RNA stained with Syto9 (green channel, left column) and surface-associated amyloid fibers stained with Congo Red (red channel, middle column). Combined channel view in right column. Scale bars represent 10 µm.

**Supplementary Figure 5.**
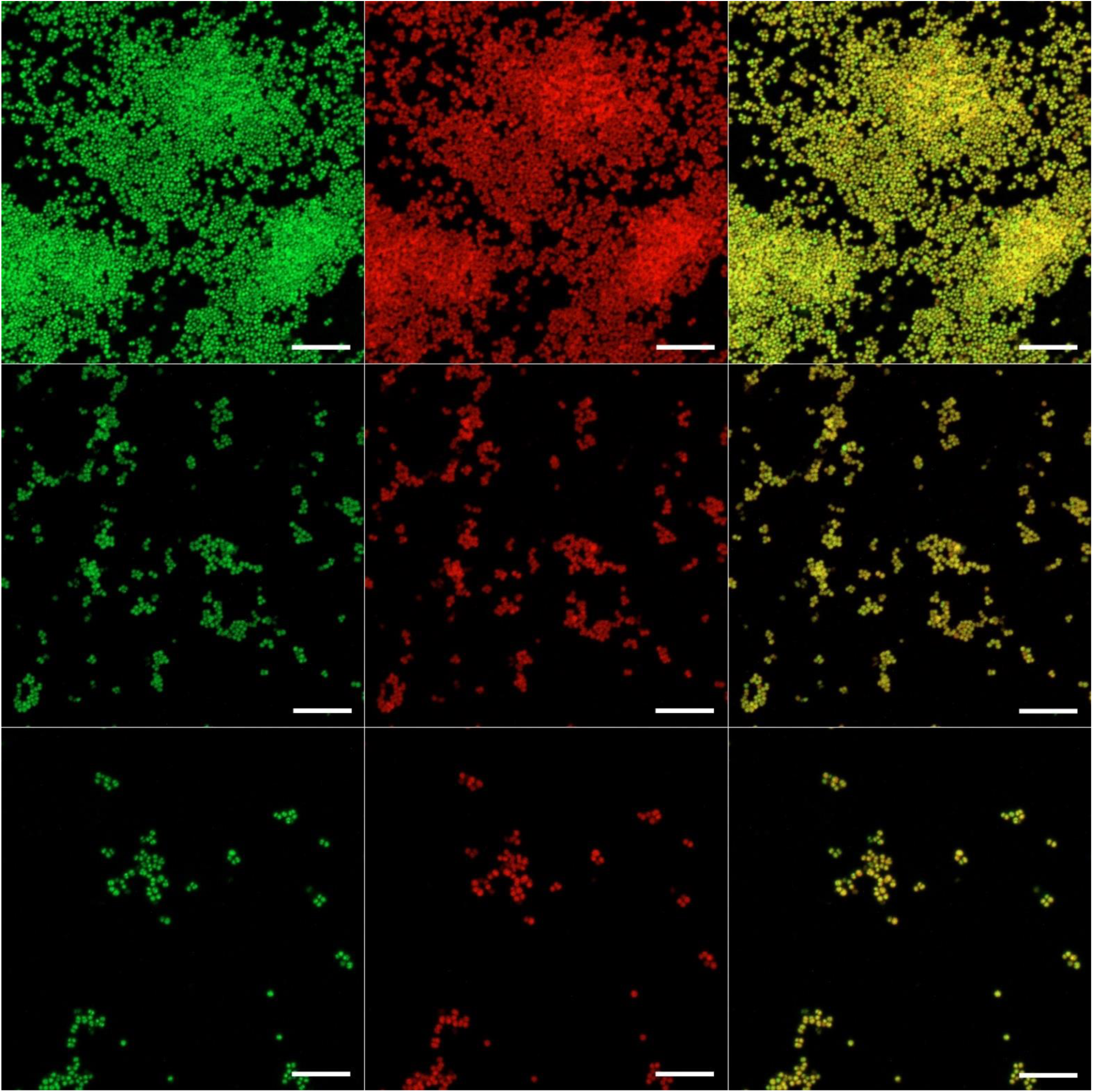
Maximal orthogonal projections of representative CLSM images of fixed 48 h growth medium *S. aureus* biofilm on uncoated surface (upper panel), sparse nano-ZnO surface (middle panel) and dense nano-ZnO (bottom panel). DNA/RNA stained with Syto9 (green channel, left column) and surface-associated amyloid fibers stained with Congo Red (red channel, middle column). Combined channel view in right column. Scale bars represent 10 µm.

**Supplementary Figure 6.**
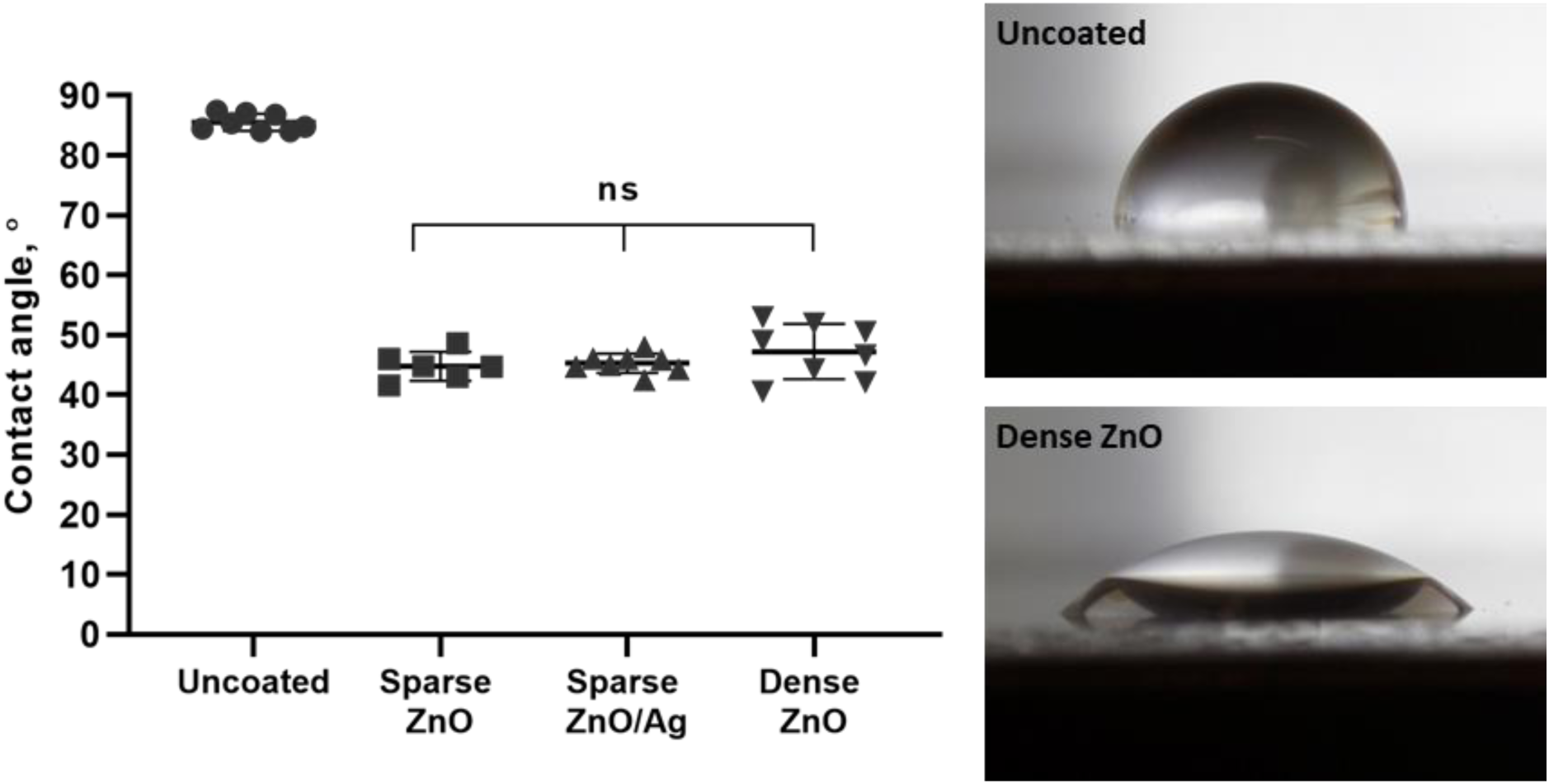
Water contact angles of uncoated, nano-ZnO and nano-ZnO/Ag surfaces. The angles were measured from photos taken 5 sec after pipetting 5 µL of water to each surface, using ImageJ software. ns – not statistically significant

**Supplementary Figure 7.**
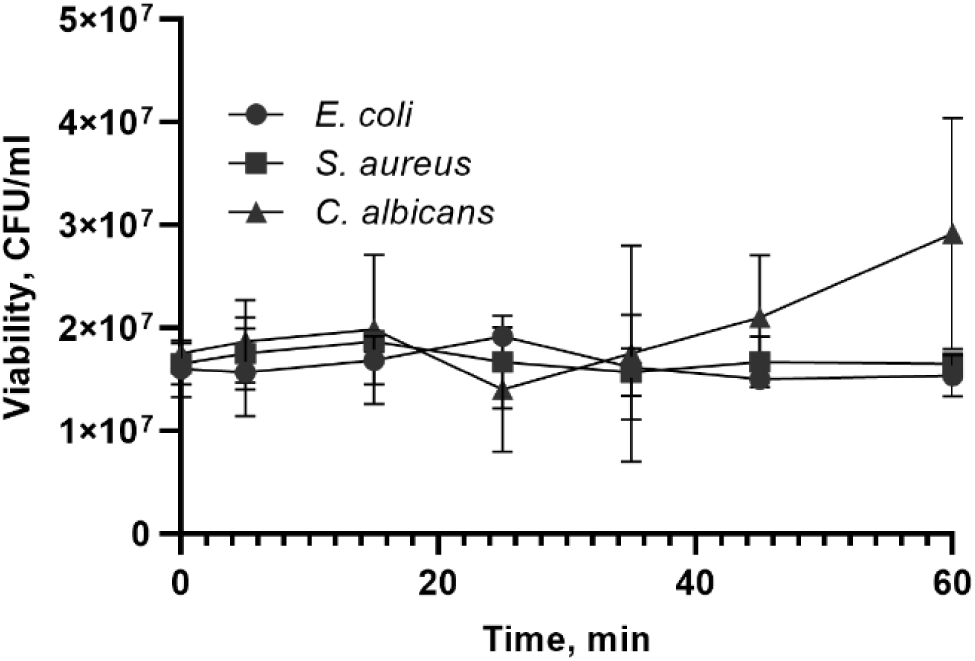
Microbial viability in biofilm harvesting medium (0.5x SCDLP with 1.5M NaCl, Table 1). 15 ml microbial was vortexed for 30 sec, sonicated for 0-60 min, vortexed again for 30 sec, serially diluted and plated for counting. Harvesting protocol in 0.5x SCDLP with high salt concentration did not significantly affect microbial viability after 15 min sonication compared to the initial suspension (P≥0.05).

**Supplementary Table 1.** Zn and Ag content in the nano-ZnO and nano-ZnO/Ag surfaces; theoretical maximum release into 5 ml microbial test medium. ND denotes “not detected”.

| Surface coating | Zn<br>$\mu\text{g}/\text{cm}^2$ | SD | Max Zn<br>release<br>into 5 ml,<br>$\mu\text{g}/\text{ml}$ | SD | Ag<br>$\text{ng}/\text{cm}^2$ | SD | Max Ag<br>release<br>into 5 ml,<br>$\text{ng}/\text{ml}$ | SD |
| --- | --- | --- | --- | --- | --- | --- | --- | --- |
| Uncoated | 0.02 | 0.00 | 0.01 | 0.00 | ND | ND | ND | ND |
| Sparse nano-ZnO | 2.12 | 0.12 | 1.36 | 0.08 | ND | ND | ND | ND |
| Sparse nano-ZnO/Ag | 2.39 | 0.10 | 1.54 | 0.07 | 23.56 | 3.42 | 15.27 | 2.21 |
| Dense nano-ZnO | 18.00 | 0.70 | 11.65 | 0.45 | ND | ND | ND | ND |

## REFERENCES

[1] H.-C. Flemming and S. Wuertz, “Bacteria and archaea on Earth and their abundance in biofilms,” Nature Reviews Microbiology, vol. 17, no. 4, pp. 247–260, Apr. 2019, doi: 10.1038/s41579-019-0158-9.

[2] NIH, “Research on microbial biofilms: PA Number: PA-03-047,” National Institute of Health, Dec. 2002. [Online]. Available: http://grants.nih.gov/grants/guide/pa-files/PA-03-047.html.

[3] J. N. Pendleton, S. P. Gorman, and B. F. Gilmore, “Clinical relevance of the ESKAPE pathogens,” Expert Review of Anti-infective Therapy, vol. 11, no. 3, pp. 297–308, Mar. 2013, doi: 10.1586/eri.13.12.

[4] H. Ceri, M. E. Olson, C. Stremick, R. R. Read, D. Morck, and A. Buret, “The Calgary Biofilm Device: new technology for rapid determination of antibiotic susceptibilities of bacterial biofilms,” J. Clin. Microbiol., vol. 37, no. 6, pp. 1771–1776, Jun. 1999.

[5] M. T. France, A. Cornea, H. Kehlet-Delgado, and L. J. Forney, “Spatial structure facilitates the accumulation and persistence of antibiotic-resistant mutants in biofilms,” Evol Appl, vol. 12, no. 3, pp. 498–507, Mar. 2019, doi: 10.1111/eva.12728.

[6] I. Olsen, “Biofilm-specific antibiotic tolerance and resistance,” Eur J Clin Microbiol Infect Dis, vol. 34, no. 5, pp. 877–886, May 2015, doi: 10.1007/s10096-015-2323-z.

[7] A. Santos-Lopez, C. W. Marshall, M. R. Scribner, D. J. Snyder, and V. S. Cooper, “Evolutionary pathways to antibiotic resistance are dependent upon environmental structure and bacterial lifestyle,” Elife, vol. 8, 13 2019, doi: 10.7554/eLife.47612.

[8] J. Sjollema et al., “In vitro methods for the evaluation of antimicrobial surface designs,” Acta Biomaterialia, vol. 70, pp. 12–24, Apr. 2018, doi: 10.1016/j.actbio.2018.02.001.

[9] M. Rosenberg, N. F. Azevedo, and A. Ivask, “Propidium iodide staining underestimates viability of adherent bacterial cells,” Scientific Reports, vol. 9, no. 1, Dec. 2019, doi: 10.1038/s41598-019-42906-3.

[10] K. N. Kragh, M. Alhede, L. Kvich, and T. Bjarnsholt, “Into the well—A close look at the complex structures of a microtiter biofilm and the crystal violet assay,” Biofilm, p. 100006, Sep. 2019, doi: 10.1016/j.bioflm.2019.100006.

[11] K. D. Mandakhalikar, J. N. Rahmat, E. Chiong, K. G. Neoh, L. Shen, and P. A. Tambyah, “Extraction and quantification of biofilm bacteria: Method optimized for urinary catheters,” Sci Rep, vol. 8, no. 1, p. 8069, Dec. 2018, doi: 10.1038/s41598-018-26342-3.

[12] H. Kobayashi, M. Oethinger, M. J. Tuohy, G. W. Procop, and T. W. Bauer, “Improved detection of biofilm-formative bacteria by vortexing and sonication: a pilot study,” Clin. Orthop. Relat. Res., vol. 467, no. 5, pp. 1360–1364, May 2009, doi: 10.1007/s11999-008-0609-5.

[13] J. J. Harrison, H. Ceri, C. A. Stremick, and R. J. Turner, “Biofilm susceptibility to metal toxicity,” Environ Microbiol, vol. 6, no. 12, pp. 1220–1227, Dec. 2004, doi: 10.1111/j.1462-2920.2004.00656.x.

[14] J. A. Lemire, J. J. Harrison, and R. J. Turner, “Antimicrobial activity of metals: mechanisms, molecular targets and applications,” Nat Rev Microbiol, vol. 11, no. 6, pp. 371–384, Jun. 2013, doi: 10.1038/nrmicro3028.

[15] C. Andreini, L. Banci, I. Bertini, and A. Rosato, “Zinc through the Three Domains of Life,” J. Proteome Res., vol. 5, no. 11, pp. 3173–3178, Nov. 2006, doi: 10.1021/pr0603699.

[16] J. J. Harrison et al., “Chromosomal antioxidant genes have metal ion-specific roles as determinants of bacterial metal tolerance,” Environmental Microbiology, vol. 11, no. 10, pp. 2491–2509, Oct. 2009, doi: 10.1111/j.1462-2920.2009.01973.x.

[17] R. M. Couñago et al., “Imperfect coordination chemistry facilitates metal ion release in the Psa permease,” Nat Chem Biol, vol. 10, no. 1, pp. 35–41, Jan. 2014, doi: 10.1038/nchembio.1382.

[18] C. A. McDevitt et al., “A Molecular Mechanism for Bacterial Susceptibility to Zinc,” PLoS Pathog, vol. 7, no. 11, p. e1002357, Nov. 2011, doi: 10.1371/journal.ppat.1002357.

[19] C. Y. Ong, M. J. Walker, and A. G. McEwan, “Zinc disrupts central carbon metabolism and capsule biosynthesis in Streptococcus pyogenes,” Sci Rep, vol. 5, no. 1, p. 10799, Sep. 2015, doi: 10.1038/srep10799.

[20] D. A. Mills, B. Schmidt, C. Hiser, E. Westley, and S. Ferguson-Miller, “Membrane Potential-controlled Inhibition of Cytochrome *c* Oxidase by Zinc,” J. Biol. Chem., vol. 277, no. 17, pp. 14894–14901, Apr. 2002, doi: 10.1074/jbc.M111922200.

[21] J. P. Hosler, S. Ferguson-Miller, and D. A. Mills, “Energy Transduction: Proton Transfer Through the Respiratory Complexes,” Annu. Rev. Biochem., vol. 75, no. 1, pp. 165–187, Jun. 2006, doi: 10.1146/annurev.biochem.75.062003.101730.

[22] B. A. Eijkelkamp et al., “Extracellular Zinc Competitively Inhibits Manganese Uptake and Compromises Oxidative Stress Management in Streptococcus pneumoniae,” PLoS ONE, vol. 9, no. 2, p. e89427, Feb. 2014, doi: 10.1371/journal.pone.0089427.

[23] J. Pasquet, Y. Chevalier, J. Pelletier, E. Couval, D. Bouvier, and M.-A. Bolzinger, “The contribution of zinc ions to the antimicrobial activity of zinc oxide,” Colloids and Surfaces A: Physicochemical and Engineering Aspects, vol. 457, pp. 263–274, Sep. 2014, doi: 10.1016/j.colsurfa.2014.05.057.

[24] A. Sirelkhatim et al., “Review on Zinc Oxide Nanoparticles: Antibacterial Activity and Toxicity Mechanism,” Nano-Micro Lett., vol. 7, no. 3, pp. 219–242, Jul. 2015, doi: 10.1007/s40820-015-0040-x.

[25] J. T. Seil and T. J. Webster, “Reduced Staphylococcus aureus proliferation and biofilm formation on zinc oxide nanoparticle PVC composite surfaces,” Acta Biomaterialia, vol. 7, no. 6, pp. 2579–2584, Jun. 2011, doi: 10.1016/j.actbio.2011.03.018.

[26] R. Pati et al., “Topical application of zinc oxide nanoparticles reduces bacterial skin infection in mice and exhibits antibacterial activity by inducing oxidative stress response and cell membrane disintegration in macrophages,” Nanomedicine: Nanotechnology, Biology and Medicine, vol. 10, no. 6, pp. 1195–1208, Aug. 2014, doi: 10.1016/j.nano.2014.02.012.

[27] J.-H. Lee, Y.-G. Kim, M. H. Cho, and J. Lee, “ZnO nanoparticles inhibit Pseudomonas aeruginosa biofilm formation and virulence factor production,” Microbiological Research, vol. 169, no. 12, pp. 888–896, Dec. 2014, doi: 10.1016/j.micres.2014.05.005.

[28] L. J. Lee, J. A. Barrett, and R. K. Poole, “Genome-wide transcriptional response of chemostat-cultured Escherichia coli to zinc,” J. Bacteriol., vol. 187, no. 3, pp. 1124–1134, Feb. 2005, doi: 10.1128/JB.187.3.1124-1134.2005.

[29] R. Brayner, R. Ferrari-Iliou, N. Brivois, S. Djediat, M. F. Benedetti, and F. Fiévet, “Toxicological Impact Studies Based on *Escherichia coli* Bacteria in Ultrafine ZnO Nanoparticles Colloidal Medium,” Nano Lett., vol. 6, no. 4, pp. 866–870, Apr. 2006, doi: 10.1021/nl052326h.

[30] R. K. Dutta, B. P. Nenavathu, M. K. Gangishetty, and A. V. R. Reddy, “Studies on antibacterial activity of ZnO nanoparticles by ROS induced lipid peroxidation,” Colloids and Surfaces B: Biointerfaces, vol. 94, pp. 143–150, Jun. 2012, doi: 10.1016/j.colsurfb.2012.01.046.

[31] P. D. Bragg and D. J. Rainnie, “The effect of silver ions on the respiratory chain of *Escherichia coli*,” Can. J. Microbiol., vol. 20, no. 6, pp. 883–889, Jun. 1974, doi: 10.1139/m74-135.

[32] H.-J. Park et al., “Silver-ion-mediated reactive oxygen species generation affecting bactericidal activity,” Water Research, vol. 43, no. 4, pp. 1027–1032, Mar. 2009, doi: 10.1016/j.watres.2008.12.002.

[33] S. Y. Liau, D. C. Read, W. J. Pugh, J. R. Furr, and A. D. Russell, “Interaction of silver nitrate with readily identifiable groups: relationship to the antibacterialaction of silver ions,” Letters in Applied Microbiology, vol. 25, no. 4, pp. 279–283, Sep. 1997, doi: 10.1046/j.1472-765X.1997.00219.x.

[34] C.-N. Lok et al., “Silver nanoparticles: partial oxidation and antibacterial activities,” J Biol Inorg Chem, vol. 12, no. 4, pp. 527–534, Apr. 2007, doi: 10.1007/s00775-007-0208-z.

[35] A. D. Russell and W. B. Hugo, “7 Antimicrobial Activity and Action of Silver,” in Progress in Medicinal Chemistry, vol. 31, Elsevier, 1994, pp. 351–370.

[36] W. K. Jung, H. C. Koo, K. W. Kim, S. Shin, S. H. Kim, and Y. H. Park, “Antibacterial Activity and Mechanism of Action of the Silver Ion in Staphylococcus aureus and Escherichia coli,” AEM, vol. 74, no. 7, pp. 2171–2178, Apr. 2008, doi: 10.1128/AEM.02001-07.

[37] C.-N. Lok et al., “Proteomic Analysis of the Mode of Antibacterial Action of Silver Nanoparticles,” J. Proteome Res., vol. 5, no. 4, pp. 916–924, Apr. 2006, doi: 10.1021/pr0504079.

[38] Q. L. Feng, J. Wu, G. Q. Chen, F. Z. Cui, T. N. Kim, and J. O. Kim, “A mechanistic study of the antibacterial effect of silver ions on Escherichia coli and Staphylococcus aureus,” J. Biomed. Mater. Res., vol. 52, no. 4, pp. 662–668, Dec. 2000, doi: 10.1002/1097-4636(20001215)52:4<662::aid-jbm10>3.0.co;2-3.

[39] J. L. Clement and P. S. Jarrett, “Antibacterial Silver,” Metal-Based Drugs, vol. 1, no. 5–6, pp. 467–482, 1994, doi: 10.1155/MBD.1994.467.

[40] B. Gómez-Gómez, L. Arregui, S. Serrano, A. Santos, T. Pérez-Corona, and Y. Madrid, “Unravelling mechanisms of bacterial quorum sensing disruption by metal-based nanoparticles,” Science of The Total Environment, vol. 696, p. 133869, Dec. 2019, doi: 10.1016/j.scitotenv.2019.133869.

[41] N. A. Al-Shabib et al., “Biogenic synthesis of Zinc oxide nanostructures from Nigella sativa seed: Prospective role as food packaging material inhibiting broad-spectrum quorum sensing and biofilm,” Sci Rep, vol. 6, no. 1, p. 36761, Dec. 2016, doi: 10.1038/srep36761.

[42] B. García-Lara et al., “Inhibition of quorum-sensing-dependent virulence factors and biofilm formation of clinical and environmental *Pseudomonas aeruginosa* strains by ZnO nanoparticles,” Lett Appl Microbiol, vol. 61, no. 3, pp. 299–305, Sep. 2015, doi: 10.1111/lam.12456.

[43] F. Zähringer, E. Lacanna, U. Jenal, T. Schirmer, and A. Boehm, “Structure and Signaling Mechanism of a Zinc-Sensory Diguanylate Cyclase,” Structure, vol. 21, no. 7, pp. 1149–1157, Jul. 2013, doi: 10.1016/j.str.2013.04.026.

[44] Z. Huma et al., “Nanosilver Mitigates Biofilm Formation via FapC Amyloidosis Inhibition,” Small, p. 1906674, Jan. 2020, doi: 10.1002/smll.201906674.

[45] A. E. Yarawsky, S. L. Johns, P. Schuck, and A. B. Herr, “The biofilm adhesion protein Aap from *Staphylococcus epidermidis* forms zinc-dependent amyloid fibers,” J. Biol. Chem., vol. 295, no. 14, pp. 4411–4427, Apr. 2020, doi: 10.1074/jbc.RA119.010874.

[46] U. Joost et al., “Photocatalytic antibacterial activity of nano-TiO2 (anatase)-based thin films: Effects on Escherichia coli cells and fatty acids,” Journal of Photochemistry and Photobiology B: Biology, vol. 142, pp. 178–185, Jan. 2015, doi: 10.1016/j.jphotobiol.2014.12.010.

[47] V. Lakshmi Prasanna and R. Vijayaraghavan, “Insight into the Mechanism of Antibacterial Activity of ZnO: Surface Defects Mediated Reactive Oxygen Species Even in the Dark,” Langmuir, vol. 31, no. 33, pp. 9155–9162, Aug. 2015, doi: 10.1021/acs.langmuir.5b02266.

[48] H. Sun et al., “Zinc oxide/vanadium pentoxide heterostructures with enhanced day-night antibacterial activities,” Journal of Colloid and Interface Science, vol. 547, pp. 40–49, Jul. 2019, doi: 10.1016/j.jcis.2019.03.061.

[49] K. Hirota, M. Sugimoto, M. Kato, K. Tsukagoshi, T. Tanigawa, and H. Sugimoto, “Preparation of zinc oxide ceramics with a sustainable antibacterial activity under dark conditions,” Ceramics International, vol. 36, no. 2, pp. 497–506, Mar. 2010, doi: 10.1016/j.ceramint.2009.09.026.

[50] A. Lipovsky, Y. Nitzan, A. Gedanken, and R. Lubart, “Antifungal activity of ZnO nanoparticles—the role of ROS mediated cell injury,” Nanotechnology, vol. 22, no. 10, p. 105101, Mar. 2011, doi: 10.1088/0957-4484/22/10/105101.

[51] E. H. Abdulkareem, K. Memarzadeh, R. P. Allaker, J. Huang, J. Pratten, and D. Spratt, “Anti-biofilm activity of zinc oxide and hydroxyapatite nanoparticles as dental implant coating materials,” Journal of Dentistry, vol. 43, no. 12, pp. 1462–1469, Dec. 2015, doi: 10.1016/j.jdent.2015.10.010.

[52] P. Carvalho et al., “Influence of thickness and coatings morphology in the antimicrobial performance of zinc oxide coatings,” Applied Surface Science, vol. 307, pp. 548–557, Jul. 2014, doi: 10.1016/j.apsusc.2014.04.072.

[53] M. Visnapuu et al., “UVA-induced antimicrobial activity of ZnO/Ag nanocomposite covered surfaces,” Colloids and Surfaces B: Biointerfaces, vol. 169, pp. 222–232, Sep. 2018, doi: 10.1016/j.colsurfb.2018.05.009.

[54] International Organization for Standardization, “ISO 22196:2011 Measurement of antibacterial activity on plastics and other non-porous surfaces.” International Organization for Standardization, 2011, [Online]. Available: www.iso.org.

[55] International Organization for Standardization, “ISO 27447:2009 Fine ceramics (advanced ceramics, advanced technical ceramics) - Test method for antibacterial activity of semiconducting photocatalytic materials.” International Organization for Standardization, 2009, [Online]. Available: www.iso.org.

[56] G. A. O’Toole and R. Kolter, “Initiation of biofilm formation in *Pseudomonas fluorescens* WCS365 proceeds via multiple, convergent signalling pathways: a genetic analysis,” Molecular Microbiology, vol. 28, no. 3, pp. 449–461, Apr. 1998, doi: 10.1046/j.1365-2958.1998.00797.x.

[57] R. Herrmann, F. J. García-García, and A. Reller, “Rapid degradation of zinc oxide nanoparticles by phosphate ions,” Beilstein J. Nanotechnol., vol. 5, pp. 2007–2015, Nov. 2014, doi: 10.3762/bjnano.5.209.

[58] M. Li, L. Zhu, and D. Lin, “Toxicity of ZnO nanoparticles to Escherichia coli: mechanism and the influence of medium components,” Environ. Sci. Technol., vol. 45, no. 5, pp. 1977–1983, Mar. 2011, doi: 10.1021/es102624t.

[59] R. Behra et al., “Bioavailability of silver nanoparticles and ions: from a chemical and biochemical perspective,” J R Soc Interface, vol. 10, no. 87, p. 20130396, Oct. 2013, doi: 10.1098/rsif.2013.0396.

[60] Z.-M. Xiu, J. Ma, and P. J. J. Alvarez, “Differential effect of common ligands and molecular oxygen on antimicrobial activity of silver nanoparticles versus silver ions,” Environ. Sci. Technol., vol. 45, no. 20, pp. 9003–9008, Oct. 2011, doi: 10.1021/es201918f.

[61] S. Suppi et al., “A novel method for comparison of biocidal properties of nanomaterials to bacteria, yeasts and algae,” Journal of Hazardous Materials, vol. 286, pp. 75–84, Apr. 2015, doi: 10.1016/j.jhazmat.2014.12.027.

[62] L. Brown, J. M. Wolf, R. Prados-Rosales, and A. Casadevall, “Through the wall: extracellular vesicles in Gram-positive bacteria, mycobacteria and fungi,” Nat Rev Microbiol, vol. 13, no. 10, pp. 620–630, Oct. 2015, doi: 10.1038/nrmicro3480.

[63] E.-Y. Lee et al., “Gram-positive bacteria produce membrane vesicles: Proteomics-based characterization of Staphylococcus aureus-derived membrane vesicles,” Proteomics, vol. 9, no. 24, pp. 5425–5436, Dec. 2009, doi: 10.1002/pmic.200900338.

[64] F. Andreoni et al., “Antibiotics Stimulate Formation of Vesicles in *Staphylococcus aureus* in both Phage-Dependent and -Independent Fashions and via Different Routes,” Antimicrob Agents Chemother, vol. 63, no. 2, pp. e01439–18, Dec. 2018, doi: 10.1128/AAC.01439-18.

[65] C. Formosa-Dague, P. Speziale, T. J. Foster, J. A. Geoghegan, and Y. F. Dufrêne, “Zinc-dependent mechanical properties of Staphylococcus aureus biofilm-forming surface protein SasG,” Proc. Natl. Acad. Sci. U.S.A., vol. 113, no. 2, pp. 410–415, Jan. 2016, doi: 10.1073/pnas.1519265113.

[66] D. G. Conrady, C. C. Brescia, K. Horii, A. A. Weiss, D. J. Hassett, and A. B. Herr, “A zinc-dependent adhesion module is responsible for intercellular adhesion in staphylococcal biofilms,” Proc. Natl. Acad. Sci. U.S.A., vol. 105, no. 49, pp. 19456–19461, Dec. 2008, doi: 10.1073/pnas.0807717105.

[67] P. Sudbery, N. Gow, and J. Berman, “The distinct morphogenic states of Candida albicans,” Trends in Microbiology, vol. 12, no. 7, pp. 317–324, Jul. 2004, doi: 10.1016/j.tim.2004.05.008.

[68] H.-C. Flemming and J. Wingender, “The biofilm matrix,” Nature Reviews Microbiology, Aug. 2010, doi: 10.1038/nrmicro2415.

[69] P. Stiefel, S. Schmidt-Emrich, K. Maniura-Weber, and Q. Ren, “Critical aspects of using bacterial cell viability assays with the fluorophores SYTO9 and propidium iodide,” BMC Microbiology, vol. 15, no. 1, p. 36, 2015, doi: 10.1186/s12866-015-0376-x.

[70] N. Gugala, J. A. Lemire, and R. J. Turner, “The efficacy of different anti-microbial metals at preventing the formation of, and eradicating bacterial biofilms of pathogenic indicator strains,” J Antibiot, vol. 70, no. 6, pp. 775–780, Jun. 2017, doi: 10.1038/ja.2017.10.

[71] K. Schwartz, A. K. Syed, R. E. Stephenson, A. H. Rickard, and B. R. Boles, “Functional amyloids composed of phenol soluble modulins stabilize Staphylococcus aureus biofilms,” PLoS Pathog., vol. 8, no. 6, p. e1002744, 2012, doi: 10.1371/journal.ppat.1002744.

[72] M. M. Barnhart and M. R. Chapman, “Curli Biogenesis and Function,” Annual Review of Microbiology, vol. 60, no. 1, pp. 131–147, Oct. 2006, doi: 10.1146/annurev.micro.60.080805.142106.

[73] A. Taglialegna et al., “Staphylococcal Bap Proteins Build Amyloid Scaffold Biofilm Matrices in Response to Environmental Signals,” PLoS Pathog., vol. 12, no. 6, p. e1005711, 2016, doi: 10.1371/journal.ppat.1005711.

[74] A. Dutta, S. Bhattacharyya, A. Kundu, D. Dutta, and A. K. Das, “Macroscopic amyloid fiber formation by staphylococcal biofilm associated SuhB protein,” Biophys. Chem., vol. 217, pp. 32–41, 2016, doi: 10.1016/j.bpc.2016.07.006.

[75] V. Tõugu et al., “Zn(II)- and Cu(II)-induced non-fibrillar aggregates of amyloid-β (1-42) peptide are transformed to amyloid fibrils, both spontaneously and under the influence of metal chelators,” Journal of Neurochemistry, vol. 110, no. 6, pp. 1784–1795, Sep. 2009, doi: 10.1111/j.1471-4159.2009.06269.x.

[76] C. J. Sarell, S. R. Wilkinson, and J. H. Viles, “Substoichiometric Levels of Cu ^2+^ Ions Accelerate the Kinetics of Fiber Formation and Promote Cell Toxicity of Amyloid-β from Alzheimer Disease,” Journal of Biological Chemistry, vol. 285, no. 53, pp. 41533–41540, Dec. 2010, doi: 10.1074/jbc.M110.171355.

[77] A. Abelein, A. Gräslund, and J. Danielsson, “Zinc as chaperone-mimicking agent for retardation of amyloid β peptide fibril formation,” Proceedings of the National Academy of Sciences, vol. 112, no. 17, pp. 5407–5412, Apr. 2015, doi: 10.1073/pnas.1421961112.

[78] B. Ma, F. Zhang, X. Wang, and X. Zhu, “Investigating the inhibitory effects of zinc ions on amyloid fibril formation of hen egg-white lysozyme,” International Journal of Biological Macromolecules, vol. 98, pp. 717–722, May 2017, doi: 10.1016/j.ijbiomac.2017.01.128.

[79] D. K. Ban and S. Paul, “Nano Zinc Oxide Inhibits Fibrillar Growth and Suppresses Cellular Toxicity of Lysozyme Amyloid,” ACS Applied Materials & Interfaces, vol. 8, no. 46, pp. 31587–31601, Nov. 2016, doi: 10.1021/acsami.6b11549.

[80] D. Wilson, “Candida albicans,” Trends in Microbiology, vol. 27, no. 2, pp. 188–189, Feb. 2019, doi: 10.1016/j.tim.2018.10.010.

[81] S. Kurakado, R. Arai, and T. Sugita, “Association of the hypha-related protein Pra1 and zinc transporter Zrt1 with biofilm formation by the pathogenic yeast *Candida albicans*: Pra1 and Zrt1 regulate biofilm formation,” Microbiol Immunol, vol. 62, no. 6, pp. 405–410, Jun. 2018, doi: 10.1111/1348-0421.12596.

[82] M. Cierech et al., “Significance of polymethylmethacrylate (PMMA) modification by zinc oxide nanoparticles for fungal biofilm formation,” International Journal of Pharmaceutics, vol. 510, no. 1, pp. 323–335, Aug. 2016, doi: 10.1016/j.ijpharm.2016.06.052.

[83] M. Jalal, M. A. Ansari, S. G. Ali, H. M. Khan, and S. Rehman, “Anticandidal activity of bioinspired ZnO NPs: effect on growth, cell morphology and key virulence attributes of *Candida* species,” Artificial Cells, Nanomedicine, and Biotechnology, vol. 46, no. sup1, pp. 912–925, Oct. 2018, doi: 10.1080/21691401.2018.1439837.

[84] M. Divya et al., “Biopolymer gelatin-coated zinc oxide nanoparticles showed high antibacterial, antibiofilm and anti-angiogenic activity,” Journal of Photochemistry and Photobiology B: Biology, vol. 178, pp. 211–218, Jan. 2018, doi: 10.1016/j.jphotobiol.2017.11.008.

[85] J. J. Harrison et al., “Metal Ions May Suppress or Enhance Cellular Differentiation in Candida albicans and Candida tropicalis Biofilms,” Applied and Environmental Microbiology, vol. 73, no. 15, pp. 4940–4949, Aug. 2007, doi: 10.1128/AEM.02711-06.

[86] C. J. Nobile et al., “Biofilm Matrix Regulation by Candida albicans Zap1,” PLoS Biol, vol. 7, no. 6, p. e1000133, Jun. 2009, doi: 10.1371/journal.pbio.1000133.

[87] S. Ganguly et al., “Zap1 Control of Cell-Cell Signaling in Candida albicans Biofilms,” Eukaryotic Cell, vol. 10, no. 11, pp. 1448–1454, Nov. 2011, doi: 10.1128/EC.05196-11.

[88] C. J. Nobile and A. P. Mitchell, “Regulation of Cell-Surface Genes and Biofilm Formation by the C. albicans Transcription Factor Bcr1p,” Current Biology, vol. 15, no. 12, pp. 1150–1155, Jun. 2005, doi: 10.1016/j.cub.2005.05.047.

[89] K. Forsberg et al., “*Candida auris* : The recent emergence of a multidrug-resistant fungal pathogen,” Medical Mycology, vol. 57, no. 1, pp. 1–12, Jan. 2019, doi: 10.1093/mmy/myy054.

[90] H. Wisplinghoff et al., “Nosocomial bloodstream infections due to Candida spp. in the USA: species distribution, clinical features and antifungal susceptibilities,” International Journal of Antimicrobial Agents, vol. 43, no. 1, pp. 78–81, Jan. 2014, doi: 10.1016/j.ijantimicag.2013.09.005.

[91] S. A. Klotz, B. S. Chasin, B. Powell, N. K. Gaur, and P. N. Lipke, “Polymicrobial bloodstream infections involving Candida species: analysis of patients and review of the literature,” Diagnostic Microbiology and Infectious Disease, vol. 59, no. 4, pp. 401–406, Dec. 2007, doi: 10.1016/j.diagmicrobio.2007.07.001.

[92] C. Bernard, V. Lemoine, M. A. Hoogenkamp, M. Girardot, B. P. Krom, and C. Imbert, “*Candida albicans* enhances initial biofilm growth of *Cutibacterium acnes* under aerobic conditions,” Biofouling, vol. 35, no. 3, pp. 350–360, Mar. 2019, doi: 10.1080/08927014.2019.1608966.

[93] E. P. Fox, E. S. Cowley, C. J. Nobile, N. Hartooni, D. K. Newman, and A. D. Johnson, “Anaerobic Bacteria Grow within Candida albicans Biofilms and Induce Biofilm Formation in Suspension Cultures,” Current Biology, vol. 24, no. 20, pp. 2411–2416, Oct. 2014, doi: 10.1016/j.cub.2014.08.057.

[94] B. Adam, G. S. Baillie, and L. J. Douglas, “Mixed species biofilms of Candida albicans and Staphylococcus epidermidis,” Journal of Medical Microbiology, vol. 51, no. 4, pp. 344–349, Apr. 2002, doi: 10.1099/0022-1317-51-4-344.

[95] P.-H. Lin, M. Sermersheim, H. Li, P. Lee, S. Steinberg, and J. Ma, “Zinc in Wound Healing Modulation,” Nutrients, vol. 10, no. 1, p. 16, Dec. 2017, doi: 10.3390/nu10010016.

[96] I. Tenaud, S. Leroy, N. Chebassier, and B. Dreno, “Zinc, copper and manganese enhanced keratinocyte migration through a functional modulation of keratinocyte integrins,” Exp Dermatol, vol. 9, no. 6, pp. 407–416, Dec. 2000, doi: 10.1034/j.1600-0625.2000.009006407.x.

[97] A. Sliwinska et al., “Genotoxicity and cytotoxicity of ZnO and Al 2 O 3 nanoparticles,” Toxicology Mechanisms and Methods, vol. 25, no. 3, pp. 176–183, Mar. 2015, doi: 10.3109/15376516.2015.1006509.

[98] N. K. Uzar, M. Abudayyak, N. Akcay, G. Algun, and G. Özhan, “Zinc oxide nanoparticles induced cyto- and genotoxicity in kidney epithelial cells,” Toxicology Mechanisms and Methods, vol. 25, no. 4, pp. 334–339, May 2015, doi: 10.3109/15376516.2015.1045654.

[99] A. Chiba, S. Sugimoto, F. Sato, S. Hori, and Y. Mizunoe, “A refined technique for extraction of extracellular matrices from bacterial biofilms and its applicability: Extraction of ECM from bacterial biofilms,” Microbial Biotechnology, vol. 8, no. 3, pp. 392–403, May 2015, doi: 10.1111/1751-7915.12155.

[100] M. Slifkin and R. Cumbie, “Congo red as a fluorochrome for the rapid detection of fungi,” J. Clin. Microbiol., vol. 26, no. 5, pp. 827–830, May 1988.

[101] P. N. Lipke, M. C. Garcia, D. Alsteens, C. B. Ramsook, S. A. Klotz, and Y. F. Dufrêne, “Strengthening relationships: amyloids create adhesion nanodomains in yeasts,” Trends in Microbiology, vol. 20, no. 2, pp. 59–65, Feb. 2012, doi: 10.1016/j.tim.2011.10.002.

[102] D. O. Serra, A. M. Richter, and R. Hengge, “Cellulose as an Architectural Element in Spatially Structured Escherichia coli Biofilms,” Journal of Bacteriology, vol. 195, no. 24, pp. 5540–5554, Dec. 2013, doi: 10.1128/JB.00946-13.

